# The structural basis for *de novo* DNA methylation in chromatin

**DOI:** 10.1101/2024.12.19.629503

**Authors:** Xiaoyan Xie, Minmin Liu, X. Edward Zhou, Michelle L. Dykstra, Peter A. Jones, Evan J. Worden

## Abstract

*De novo* cytosine methylation is essential for mammalian development and is deposited by DNMT3A and DNMT3B. In cells, DNA methylation occurs in the context of chromatin, where nucleosomes are connected by DNA linkers. Here, we report Cryo-EM structures of DNMT3A2/3B3 bound to di-nucleosomes with different linker lengths. We show that DNMT3A2/3B3 preferentially binds di-nucleosomes separated by short DNA linkers by inducing large-scale changes to the di-nucleosome structure, enabling each DNMT3B3 subunit to bind each nucleosome. Linker length and the position of cytosines within the linker control DNA methylation, indicating that a significant fraction of linkers in chromatin are naturally resistant to DNMT3A2/3B3 activity. Finally, DNMT3A2/3B3 scans for H3K36me2-3 modifications, explaining how H3K36 methylation simulates DNMT3A2 activity. Our structure is the first example of a DNA methyltransferase interacting with higher-order nucleosome substrates and provides new insights on how DNA methylation takes place in chromatin.

## Introduction

Different patterns of methylated CpG dinucleotides (mCpG) have distinct effects on gene expression, but in general mCpG modifications at promotor regions are associated with transcriptional repression^1^. In addition, mCpG plays critical roles in establishing constitutive heterochromatin^2,3^, directing various developmental programs^4,5^ and regulating interactions between DNA and various transcription factors^6–8^. Importantly, dysregulated deposition of mCpG underlies various human diseases including cancer^9^. Therefore, proper cell physiology requires highly regulated mechanisms to direct the correct deposition of mCpG in the genome.

In humans, DNA methylation patterns are established by highly conserved *de novo* DNA methyltransferases. DNMT3A and DNMT3B are responsible for depositing the initial mCpG modifications at specific genomic loci^5^, while DNMT1 is a maintenance enzyme that recognizes hemi-methylated CpG dinucleotides and generates symmetrically methylated mCpG palindromes^10,11^. Although DNMT3A and DNMT3B are highly homologous, each enzyme has distinct methylation targets and expression profiles^11^. DNMT3A methylates major satellite repeats and determines allele-specific imprinting during gametogenesis^12^, while DNMT3B plays a dominant role in early embryonic development and silenced minor satellite repeats^12^. In addition, multiple isoforms of DNMT3A and DNMT3B have been identified, with many having specific expression profiles in tissues and disease states^11,13^. These isoforms appear to have distinct capacities to interact with chromatin and to methylate CpG^14^, indicating that specific expression of distinct DNMT3 isoforms is a crucial regulatory mechanism for controlled deposition of mCpG.

The catalytic activities of DNMT3A and DNMT3B are tightly regulated to prevent off-target DNA methylation. DNMT3A and DNMT3B have weak DNA methylation activity in isolation, that is significantly stimulated upon binding to the accessory proteins DNMT3L^15^ or DNMT3B3 (a splice variant of DNMT3B)^16,17^, which lack catalytic activity. Active DNMT3A or DNMT3B enzymes form elongated hetero-tetrameric complexes with their accessory proteins^18^ where the catalytic DNMT3A or DNMT3B is positioned at the center and the non-catalytic DNMT3L or DNMT3B3 is positioned at the ends^18,19^. The expression of DNMT3L and DNMT3B3 is controlled so each accessory factor has specific tissue or developmental roles^20^. DNMT3L is specifically expressed early in development^15^, while DNMT3B3 is expressed widely in somatic tissues^21^.

DNMT3A and DNMT3B contain specific histone recognition domains that recruit the enzymes to chromatin and further stimulate their activity. The ATRX–DNMT3–DNMT3L (ADD) domain inhibits DNA methylation by blocking DNA binding^22^. However, interactions between the ADD domain and the unmodified H3 tail near H3K4 induce a conformational change that displaces the ADD and allows DNA to bind^22,23^. The Pro-Trp-Trp-Pro (PWWP) domain binds to H3K36me2-3^24–27^ modifications and recruits the DNMT hetero-tetramer to specific chromatin loci^24^. In addition, recent studies have shown that the PWWP domain inhibits DNMT3 activity^28,29^ and that PWWP-dependent inhibition of DNMT3 enzymes can be relieved through recognition of H3K36me2 modifications in nucleosomes^29^, but not by peptides^28^. Therefore, the ADD and PWWP domains inhibit CpG methylation by DNMT3A and DNMT3B and this inhibition is relieved by recognition of H3K36me2-3 and unmodified H3K4 in the nucleosome.

While the nucleosome is important to stimulate DNMT3A and DNMT3B activity, the nucleosome DNA itself is not accessible for methylation. The histone octamer core imposes a strong barrier to DNA methylation by physically blocking access to the DNMT heterotetramer^30^. Indeed, at well-positioned nucleosomes surrounding CTCF binding sites, mCpG is preferentially deposited in linker DNA^31^. In addition, a Cryo-EM structure of the DNMT3A2/DNMT3B3 hetero-tetramer bound to a mono-nucleosome indicates that DNMT3A2/3B3 may prefer higher-order chromatin substrates that allow for multivalent interactions with DNMT3B3^19^. However, interactions between DNMT3A2/3B3 and more complex nucleosome substrates have not been investigated and it is not known how methylation of linker DNA might be regulated in the context of chromatin substrates with different nucleosome spacings.

Here we report cryo-EM structures of the DNMT3A2/3B3 hetero-tetramer bound to di-nucleosome substrates with various linker lengths in an initial binding state that is poised for activation and DNA methylation. We find that the DNMT3A2/3B3 hetero-tetramer preferentially binds to di-nucleosomes with short linkers (5-8bp) by simultaneously bridging acidic patch residues on both nucleosome surfaces. DNMT3A2/3B3 binding is accompanied by large-scale rearrangements of the di-nucleosome structure which allows each copy of DNMT3B3 to interact with each nucleosome. While di-nucleosomes with short linker lengths tightly engage the DNMT3A2/3B3 hetero-tetramer, the short DNA linkers themselves do not interact with DNMT3A2 and are not methylated. DNA linker length and the position of CpG dinucleotides relative to the nucleosome determines the extent of linker DNA methylation by DNMT3A2/3B3. Finally, we show that the ADD domains of DNMT3A2 and DNMT3B3 specifically position the PWWP domain adjacent to the H3 tail exit point where it can scan the modification state of H3K36. This configuration suggests a mechanism for catalytic activation of the DNMT3A2/3B3 hetero-tetramer and provides a structural explanation for the dual requirement of H3K36me2-3 and unmethylated H3K4 for DNA methylation.

## Results

### The DNMT3A2/3B3 hetero-tetramer binds di-nucleosomes with short linkers

We reasoned that the linker length between neighboring nucleosomes would be critical to optimally position a second nucleosome in such a way to interact with the free DNMT3B3 catalytic-like domain (**Fig. S1a**). Previous chemical mapping of nucleosome positions in mESC cells^32^ and ChIP-seq studies on human cells^33^ have shown that that the median linker length in mammalian chromatin is approximately 35bp. However, the linker length distribution is not uniform, showing peaks in linker length every 10n + 5 base pairs, with the most prominent peak at 35bp (**Fig. S1b**). This preferred nucleosome spacing is conserved across eukaryotes (**Fig S1b**)^32,34^ and is likely driven by constraints in the relative orientation between neighboring nucleosomes^35,36^. A model of the di-nucleosome containing a 35bp linker shows that this linker length places the second nucleosome too far away to engage the second DNMT3B3 subunit (**Fig. S1c**). Therefore, we generated di-nucleosome substrates with shorter linker lengths of 6bp or 25bp (which correspond to local peaks in the linker length distribution) with 10bp overhangs on each nucleosome and assessed complex formation with the DNMT3A2/3B3 hetero-tetramer using electromobility shift assays (EMSA) (**Fig. 1a**). We found that the DNMT3A2/3B3 hetero-tetramer preferentially binds di-nucleosomes separated by the 6bp linker. Reanalysis of chemically mapped nucleosome positions^32^ revealed that short-linker di-nucleosomes (0bp-10bp) make up ∼7% of all nucleosomes in mESCs (**Fig. S1d**). Therefore, nucleosomes with short linkers are a common feature of chromatin that bind tightly to DNMT3A2/3B3 *in vitro*.

**Figure 1:**
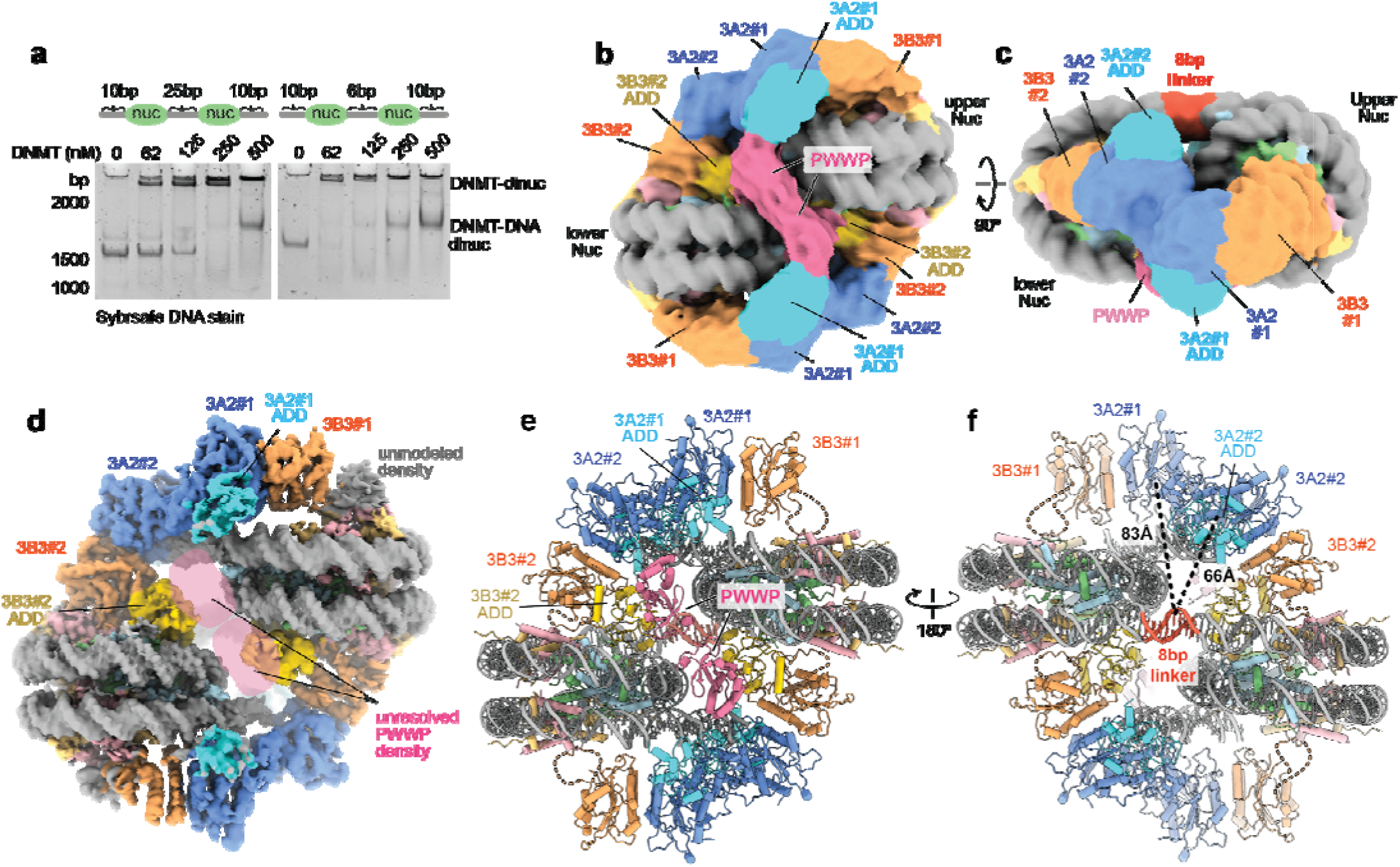
Structure of the DNMT3A2/DNMT3B3 tetramer bound to a di-nucleosome. **a** DNMT3A2/3B3 hetero-tetramers binding to di-nucleosomes with different linker lengths. Top: schematic showing di-nucleosomes with overhang and linker lengths indicated by the number of base pairs. Bottom: electromobility shift assay showing DNMT3A2/3B3 binding to di-nucleosomes. **b-c** Cryo-EM map of the complex between the DNMT3A2/3B3 tetramer and the 10-8-10 di-nucleosome filtered to 10Å and colored according to the domains of DNMT3A2, DNMT3B3 and each nucleosome. **d** Combined focused maps of each DNMT3A2 and DNMT3B3 tetramer and each nucleosome displayed in a zone around the atomic model. Unresolved density for the PWWP domains due to focused refinement is indicated with shaded regions colored pink. **e-f** Atomic model of the DNMT3A2/DNMT3B3 tetramer bound to a di-nucleosome. Protein domains are colored as in b-d. The 8bp linker DNA separating the two nucleosomes is colored red.

To understand why the DNMT3A2/3B3 hetero-tetramer prefers di-nucleosomes separated by short linkers, we determined several Cryo-EM structures (**Fig. 1b-f, S2-6, Table 1**) of the DNMT3A2/3B3 hetero-tetramer bound to unmodified di-nucleosome substrates containing 10bp overhangs and 5bp, 6bp, 8bp or 25bp linkers (hereafter referred to as 10-X-10 di-nucleosomes, where X indicates the inter-nucleosome linker length). The 1MDa complex between DNMT3A2/3B3 and the 10-8-10 di-nucleosome had better resolved density for the DNMT3A2/3B3 hetero-tetramer than other di-nucleosome complexes and was used for all subsequent analysis (**Fig. 1b-f**). The overall conformation of the DNMT3A2/3B3 hetero-tetramer on the 10-8-10 di-nucleosome is similar to the conformation of DNMT3A2/3B3 when bound to a mono-nucleosome^19^, with minor differences in the conformation of the overhang DNA (**Fig. S7a,d**). However, in this state two DNMT3A2/3B3 hetero-tetramers bind symmetrically to the di-nucleosome, which allows each DNMT3B3 subunit to interact with a nucleosome (**Fig. 1b-d**). In this conformation, the di-nucleosome adopts a stairstep-like arrangement where both nucleosome disks are aligned in parallel planes and one nucleosome is positioned a “step” up from the other. We refer to each nucleosome in the complex as either “upper” or “lower” and each subunit of the DNMT3A2/3B3 hetero-tetramer as DNMT3A2 #1 or #2 and DNMT3B3 #1 or #2 (**Fig. 1b,c**). This structure suggests that the preference of DNMT3A2/3B3 for di-nucleosomes separated by short linkers is due to contacts formed between the DNMT3A2/3B3 hetero-tetramer and both nucleosomes.

**Table 1:** Cryo-EM data collection, model refinement and validation statistics.

| Sample | DNMT3A3B/10n8n10-dinucleosome |  |  |  |  |  | 3A3B<br>10n5n10 | 10n5n10<br>alone | 3A3B<br>10n6n10 | 3A3B<br>10n8n10<br>H3<br>peptide | 3A3B<br>10n8n10<br>H2B<br>K120R | 3A3B<br>ΔPWWP<br>10n8n10 | 3A3B<br>10n25n10 | 10n8n10<br>alone |
| --- | --- | --- | --- | --- | --- | --- | --- | --- | --- | --- | --- | --- | --- | --- |
| Data Statistics |  |  |  |  |  |  |  |  |  |  |  |  |  |  |
| Microscope | Titan Krios |  |  |  |  |  | Talos<br>Arctica | Talos<br>Arctica | Titan<br>Krios | Talos<br>Arctica | Titan<br>Krios | Talos<br>Arctica | Talos<br>Arctica | Talos<br>Arctica |
| Voltage (kV) | 300 |  |  |  |  |  | 200 | 200 | 300 | 200 | 300 | 200 | 200 | 200 |
| Electron exposure (e<sup>-</sup>/Å<sup>2</sup>) | 60 |  |  |  |  |  | 50 | 50 | 60 | 50 | 60 | 50 | 50 | 50 |
| Pixel size (Å) | 0.828 |  |  |  |  |  | 0.96 | 0.96 | 0.828 | 0.96 | 0.828 | 0.96 | 0.96 | 0.96 |
| Symmetry imposed | C1 |  |  |  |  |  | C1 | C1 | C1 | C1 | C1 | C1 | C1 | C1 |
| Initial particle projections | 2584528 |  |  |  |  |  | 96074 | 367292 | 2887646 | 170287 | 331798 | 430957 | 331789 | 430957 |
| Final particle projections | 478167 |  |  |  |  |  | 40908 | 61854 | 356421 | 44359 | 103499 | 37292 | 59084 | 75131 |
| Map resolution (Å) | 3.75 |  |  |  |  |  | 6.08 | 5.93 | 3.14 | 6.08 | 3.79 | 6.92 | 4.85 | 4.24 |
| FSC threshold | 0.143 |  |  |  |  |  | 0.143 | 0.143 | 0.143 | 0.143 | 0.143 | 0.143 | 0.143 | 0.143 |
| Map resolution range (Å) | 3-10 |  |  |  |  |  | 15-5 | 10-3 | 10-3 | 15-5 | 10-3 | 15-5 | 15-5 | 10-3 |
| Model Refinement |  |  |  |  |  |  |  |  |  |  |  |  |  |  |
| EMD code | EMD-47344 | EMD-47354 | EMD-47355 | EMD-47367 | EMD-47353 | EMD-47349 | EMD-47448 | EMD-47450 | EMD-47461 | EMD-47462 | EMD-47479 | EMD-47495 | EMD-47496 | EMD-47499 |
| Structure model<br>PDB code | Composite<br>9E00 | Tetramer #1<br>9E0F | Tetramer #2<br>9E0G | Nuc 1<br>9E0R | Nuc 2<br>9E09 | Consensus<br>9E05 | 9E2D | 9E2F | 9E2Q | 9E2R | 9E3D | 9E3R | 9E3U | 9E49 |
| Model resolution (Å)<br>FSC threshold | 3.72-2.90<br>n/a | 3.46<br>0.143 | 2.90<br>0.143 | 3.20<br>0.143 | 3.72<br>0.143 | 3.75<br>0.143 | 6.08<br>0.143 | 5.93<br>0.143 | 3.14<br>0.143 | 6.08<br>0.143 | 3.79<br>0.143 | 6.92<br>0.143 | 4.85<br>0.143 | 4.24<br>0.143 |
| Model composition |  |  |  |  |  |  |  |  |  |  |  |  |  |  |
| Non-hydrogen atoms | 43875 | 12187 | 12152 | 12305 | 12355 | 49286 | 48657 | 24620 | 46712 | 49043 | 36765 | 46691 | 24818 | 24442 |
| Protein residues | 3865 | 1510 | 1543 | 749 | 752 | 4499 | 4444 | 1459 | 4188 | 4822 | 2953 | 4183 | 2248 | 1458 |
| DNA base pairs | 321 | 0 | 0 | 155 | 156 | 321 | 316 | 318 | 319 | 319 | 320 | 319 | 164 | 314 |
| Ligands | 4 | 2 | 2 | 0 | 0 | 4 | 4 | n/a | 4 | 4 | 2 | 4 | 2 | 0 |
| Zn ions | 9 | 9 | 9 | 0 | 0 | 9 | 18 | n/a | 18 | 18 | 9 | 18 | 9 | 0 |
| B factors (Å<sup>2</sup>) |  |  |  |  |  |  |  |  |  |  |  |  |  |  |
| Overall | 133.9 | 156.7 | 187.6 | 139.3 | 147.8 | 165.7 | 708.5 | 494.0 | 456.5 | 770.6 | 264.8 | 883.2 | 478.7 | 322.8 |
| Protein/DNA | 133.9 | 156.6 | 187.6 | 139.3 | 147.8 | 165.5 | 708.4 | 494.0 | 456.4 | 770.6 | 264.8 | 883.2 | 478.5 | 322.8 |
| Ligands and Zn | 310.1 | 283.5 | 287.1 | n/a | n/a | 363.3 | 912.7 | n/a | 672.3 | 770.6 | 419.7 | 1000 | 879.7 | n/a |
| R.m.s. deviations |  |  |  |  |  |  |  |  |  |  |  |  |  |  |
| Bond lengths (Å) | 0.0106 | 0.0134 | 0.0117 | 0.0054 | 0.0047 | 0.0054 | 0.0118 | 0.0083 | 0.0123 | 0.0113 | 0.0107 | 0.0116 | 0.0116 | 0.0097 |
| Bond angles (°) | 1.27 | 1.42 | 1.44 | 0.89 | 0.86 | 1.04 | 1.37 | 1.02 | 1.33 | 1.32 | 1.25 | 1.35 | 1.36 | 1.00 |
| Validation |  |  |  |  |  |  |  |  |  |  |  |  |  |  |
| MolProbity score | 1.45 | 1.61 | 1.73 | 1.07 | 1.08 | 1.33 | 1.49 | 1.15 | 1.54 | 1.53 | 1.44 | 1.55 | 1.45 | 1.09 |
| Clashscore | 5.16 | 4.46 | 4.29 | 2.81 | 2.31 | 4.15 | 4.66 | 3.21 | 5.67 | 5.44 | 4.97 | 5.68 | 4.51 | 2.98 |
| Rotamer outliers (%) | 0.0 | 0.0 | 0.0 | 0.0 | 0.0 | 0.0 | 0.03 | 0.00 | 0.00 | 0.03 | 0.08 | 0.00 | 0.00 | 0.00 |
| Ramachandran plot |  |  |  |  |  |  |  |  |  |  |  |  |  |  |
| Favored (%) | 96.95 | 96.62 | 95.53 | 98.35 | 97.68 | 97.32 | 96.26 | 97.83 | 96.47 | 96.44 | 96.92 | 96.42 | 96.59 | 98.11 |
| Allowed (%) | 3.05 | 3.38 | 4.47 | 1.65 | 2.32 | 2.68 | 3.74 | 2.17 | 3.53 | 3.56 | 3.08 | 3.58 | 3.41 | 1.89 |
| Disallowed (%) | 0 | 0 | 0 | 0 | 0 | 0 | 0 | 0 | 0 | 0 | 0 | 0 | 0 | 0 |

The ADD and PWWP domains are critical for regulating the methyltransferase activity and genomic localization of DNMT3 enzymes^22,24^. However, no structures have modeled these domains on naked DNA substrates or on nucleosomes, so it is not clear how these domains might regulate DNMT3 substrate recognition in the context of chromatin. A consensus map of the entire complex, filtered to 10Å for clarity, shows density for both DNMT3A2 catalytic domains and ADD domains (**Fig. 1b-c**). In addition, both DNMT3B3 catalytic-like domains and the ADD domain for DNMT3B3 #2 are visible, however the ADD domain from DNMT3B3 #1 is not resolved (**Fig. 1c**). The ADD domain of DNMT3B3 #2 is the most well resolved ADD in the structure (**Fig 1d**), indicating that this domain is stabilized in the context of the di-nucleosome. The map also contains a globular density positioned at the H3 tail exit site that bridges the ADD domains of DNMT3A2 #1 and DNMT3B3 #2 (**Fig. 1b**). Based on its size and position directly adjacent to the H3 tail exit site, we attribute this globular density to be the PWWP domain of either DNMT3A2 or DNMT3B3 (**Fig.1b**). We combined various focused maps to generate a single, higher-resolution composite map of the complex with density for both nucleosomes and both DNMT3A2/3B3 hetero-tetramers (**Fig. 1d**). We then used a combination of the consensus and focused maps to build a model of the entire DNMT3A2/3B3 di-nucleosome complex (**Fig. 1e,f**). Notably, the orientation of the DNMT3A2 ADD domains in our structure closely align with the ADD domains of DNMT3A1 in the autoinhibited state (**Fig S7b-c**). In this state, the ADDs do not bind the unmodified H3 tail^22^, indicating that the DNMT3A2/3B3 hetero-tetramer is autoinhibited in the structure, even though our di-nucleosome substrate contains unmodified H3. This autoinhibited state is expected to exist in solution and upon initial nucleosome engagement, prior to catalytic activation and DNA methylation, suggesting that the structure represents a pre-catalytic interaction between DNMT3A2/3B3 and the 10-8-10 di-nucleosome.

*De novo* DNA methylation at well positioned nucleosomes (e.g. near CTCF sites) is highly specific for the linker DNA between nucleosomes^37^, however the DNMT3A2/3B3 hetero-tetramer does not contact the linker DNA that separates the di-nucleosome (**Fig. 1f**). The catalytic domain of DNMT3A2 #2 binds to the 10bp overhang DNA in a similar orientation as DNMT3A2 when bound to a mono-nucleosome^19^ (**Fig. S7d**), but the catalytic active sites of DNMT3A2#1 and DNMT3A2#2 are ∼83Å and ∼66Å away from the linker DNA (**Fig. 1f**). DNMT3A2/3B3 may be prevented from accessing the linker DNA by the second nucleosome, which would clash with the DNMT hetero-tetramer when targeting the linker (**Fig. S7e-f**). This suggests that when bound to short linker di-nucleosomes, each DNMT hetero-tetramer targets the DNA protruding from either side of the di-nucleosome, not the di-nucleosome linker itself. However, in the context of chromatin, these DNA overhangs correspond to linker DNA between the di-nucleosome and neighboring nucleosomes on either side, agreeing with the preference of CpG methylation for the linker DNA between well positioned nucleosomes^37^.

### Each DNMT3B3 subunit forms distinct interactions with the nucleosome

Because the preference of DNMT3A2/3B3 for short linker di-nucleosomes likely arises from contacts formed with DNMT3B3, we examined the interactions between each DNMT3B3 subunit and the histone octamer core. A prior cryo-EM structure of the DNMT3A2/3B3 hetero-tetramer bound to a mono-nucleosome showed that the catalytic-like domain of DNMT3B3 #1 binds to the nucleosome using a pair of arginine residues (R740 and R743) that interact with the acidic patch residues H2A E56, D90, E92^19^. This contact between DNMT3B3 #1 and the upper nucleosome is recapitulated in the di-nucleosome bound structure (**Fig 2a-b**) and we observed a nearly identical contact between DNMT3B3 #2 and the lower nucleosome (**Fig 2c**). This indicates that the contacts between DNMT3B3 and the acidic patch are the same for each nucleosome in the di-nucleosome bound state. Connecting residues between the arginine anchors and the rest of DNMT3B3 were not well resolved in the structure, presumably due to high flexibility (**Fig. 2a-c**). Acidic patch interactions can occur through flexible linkers for other nucleosome binding complexes^38^, indicating that rigid connections with the acidic patch are not required for binding between the DNMT3A2/3B3 hetero-tetramer and the di-nucleosome. Additional density is also visible at each DNMT3B3 interface (**Fig. S8a**). This density is likely due to crosslinks between lysine residues in H2A and H2B as a structure of the same complex bearing an H2BK120R mutation lacks this density (**Fig. S8b**) and similar density has been observed in crosslinked nucleosome structures^39^. These data show that the DNMT3A2/3B3 hetero-tetramer uses arginine anchor residues in DNMT3B3 to engage both nucleosomes in the context of a di-nucleosome substrate.

**Figure 2:**
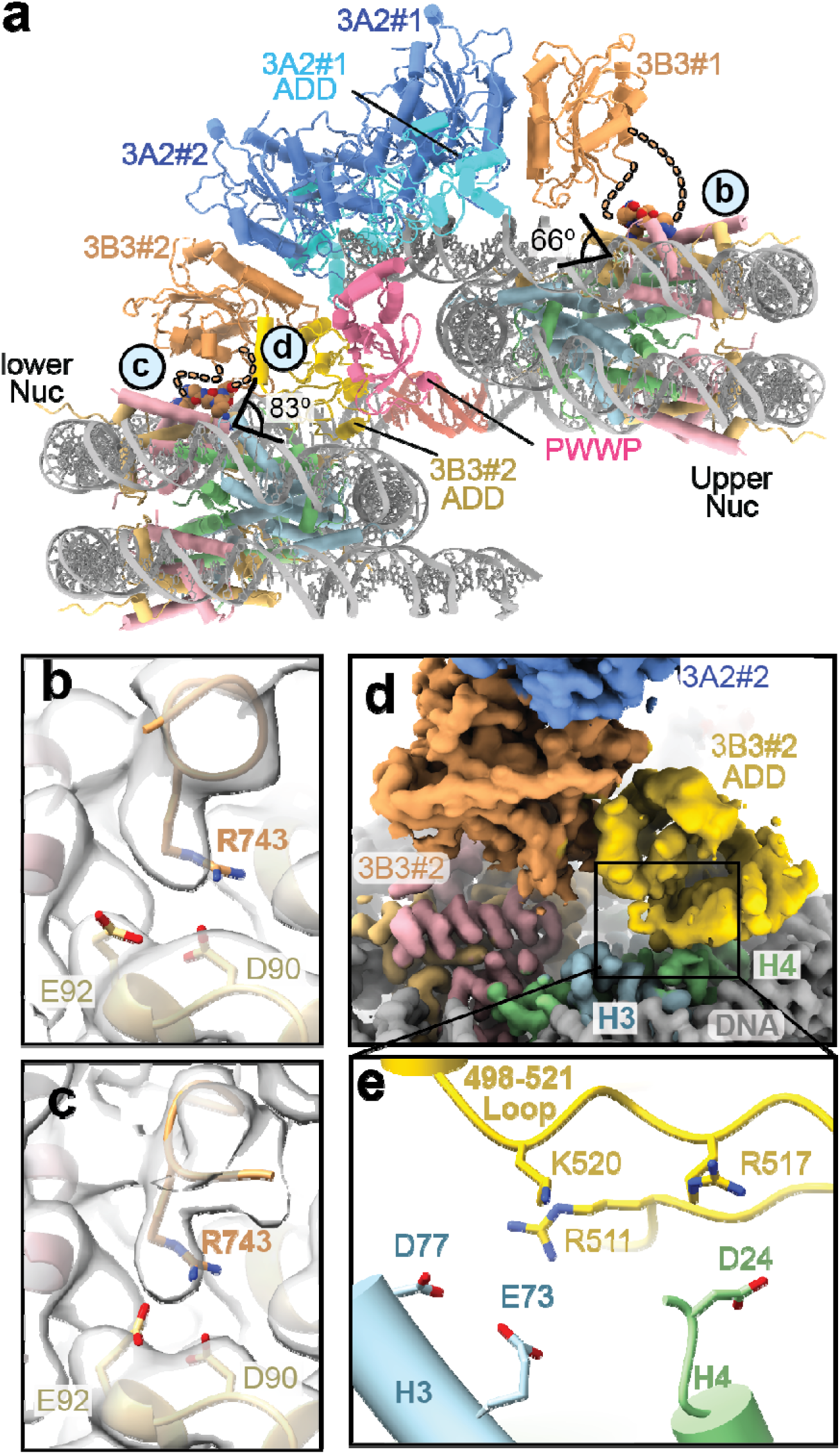
Interactions between DNMT3B3 and the histone core. **a** Overall structure of the DNMT3A2/3B3 tetramer bound to the di-nucleosome. The second copy of DNMT3A2/2/3B3 is omitted. **b-c** Close up view of acidic patch contact sites in the nucleosome focused maps. Density for the DNMT3B3 arginine anchor is shown as a semi-transparent surface. **d** Composite focused maps for the nucleosomome and DNMT3A2/3B3 tetramer to show relative resolution of each domain. **e** Close up view of the contact point betweween the DNMT3B3 #2 ADD and the histone core. Basic resides of the DNMT3B3 #2 ADD domain in proximity to acidic residues in H3 and H4 are shown in stick representation.

While residue-level interactions between the DNMT3B3 #1 and #2 catalytic-like domains and the acidic patches are similar, each DNMT3B3 subunit contacts the nucleosome in distinct conformations. The contact between DNMT3B3 #1 and the nucleosome is almost identical to the previously determined structure of the DNMT3A2/3B3 hetero-tetramer bound to a mono-nucleosome^19^. This interaction occurs at a sharp angle (∼66°) between the DNMT3B3 #1 catalytic-like domain and the nucleosome surface (**Fig. 2a**) and we observed no density for the DNMT3B3 #1 ADD (**Fig. 1c**). However, DNMT3B3 #2 binds to the lower nucleosome at an angle of 83°. This different contact angle allows the ADD from DNMT3B3 #2 to contact the nucleosome surface (**Fig. 2d**). Very diffuse density for this ADD is visible in the structure of DNMT3A2/3B3 bound to a mono-nucleosome^19^ (**Fig. S8c**), suggesting that this domain is stabilized by interacting with the lower nucleosome in a di-nucleosome substrate. The contact surface between the DNMT3B3 #2 ADD domain and the histone core is mediated through a loop in the ADD encompassing residues 498-521 that interacts with the surface of the H3/H4 dimer (**Fig. 2e**). Due to the resolution of the ADD domain, no specific side chain contacts between the ADD and the histone core can be directly inferred, and the sidechain rotamers in the model are largely determined by the crystal structure of the DNMT3B ADD domain that was used for the modeling (PDB ID: 7O45). However, the ADD 498-521 loop positions several positively charged residues (R511, R517, K520) near negatively charged residues in the H3/H4 dimer (H3 D77, H3 D73 and H4 D24) (**Fig. 2e**). This suggests that the ADD may be stabilized by electrostatic contacts with the histone surface of the lower nucleosome. The ADD from DNMT3L also has several positive charges at these positions (**Fig. S8d**), indicating that the DNMT3L ADD may interact with a nucleosome in a similar manner. Together, this shows that distinct interactions between DNMT3B3 #1, DNMT3B3 #2 and the di-nucleosome specifically stabilize the ADD domain from DNMT3B3 #2.

### Linker length dependence of di-nucleosome binding by DNMT3A2/3B3

Changes in the linker length between nucleosomes can have large effects on the relative positions of each nucleosome and affect DNMT3A2/3B3 binding (**Fig. 1a**). This is due to the periodicity of the DNA double helix, where each additional base pair in the linker rotates and separates the nucleosomes by ∼36° and ∼3.3Å. To understand how changes in the linker length affect DNMT3A2/3B3 binding, we compared Cryo-EM structures of the DNMT3A2/3B3 hetero-tetramer in complex with the 10-5-10, 10-6-10, 10-8-10 and 10-25-10 di-nucleosomes (**Fig. 3a-c, S2-6**). The DNMT-bound 10-5-10 and 10-6-10 di-nucleosomes adopt a similar conformation to the DNMT-bound 10-8-10 di-nucleosome (**Fig. 3a**). However, the lower nucleosome is rotated relative to the upper nucleosome by 22° for the 10-5-10 structure and by 8° for the 10-6-10 structure (**Fig. 3b**). Rotation of the lower nucleosome also changes the conformation of the second DNMT hetero-tetramer on the opposite face of the di-nucleosome compared to its position in the 10-8-10 complex structure (**Fig. 3b**). The lower DNMT hetero-tetramer in the 10-5-10 di-nucleosome structure is moved by 49Å and the lower DNMT hetero-tetramer in the 10-6-10 di-nucleosome moves 26Å, which agrees with the more subtle rotation of the lower nucleosome in this structure. The rotated conformation of the 10-6-10 and 10-5-10 di-nucleosomes contrast with the parallel conformation of the 10-8-10 di-nucleosome (**Fig. 1b,d**), indicating that changes in the DNA linker alter the relative conformation of the two nucleosomes in the DNMT-bound state. However, the 22° rotation of the lower nucleosome in the 10-5-10 structure (**Fig. 3b**) is less than the expected rotation of ∼100° for b-form DNA based on the 3bp difference in linker length and the periodicity of the DNA helix. Therefore, the reduced rotation of the lower nucleosome indicates that DNMT3A2/3B3 restrains the conformation of the di-nucleosome.

**Figure 3:**
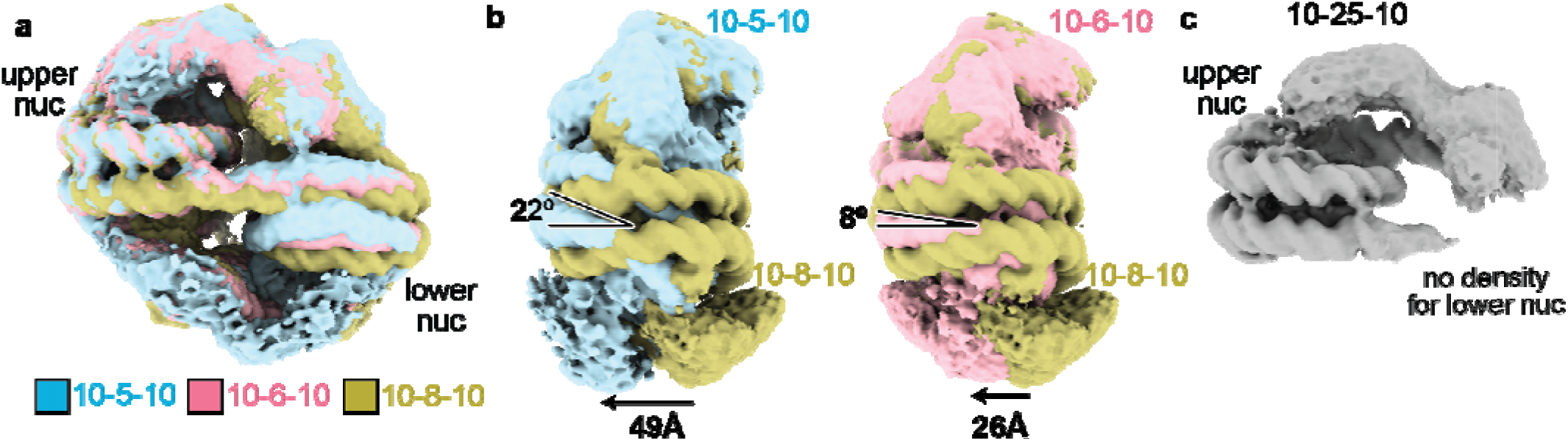
Linker length dependence of DNMT3A2/3B3 di-nucleosome bridging. **a** Superimposed consensus cryo-EM maps of the DNMT3A2/3B3 complex with 10-5-10 (blue), 10-6-10 (pink) a d 10-8-10 (yellow) di-nucleosomes. **b** Consensus cryo-EM maps of the 10-8-10 DNMT3A2/3B3 complex superimposed with the 10-5-10 (left) and 10-6-10 (right) DNMT3A2/3B3 complexes. Rotations in the lower nucleosome are indicated with the black angle indicator. **c** Consensus Cryo-EM map of the 10-25-10 DNMT3A2/3B3 complex with missing density for the lower nucleosome.

The structure of DNMT3A2/3B3 bound to the 10-25-10 di-nucleosome shows clear density for the DNMT hetero-tetramer in a conformation that matches its interaction with the “upper” nucleosome with no density for the “lower” nucleosome (**Fig. 3c**). The loss of lower nucleosome density indicates that the lower nucleosome is highly mobile. This mobility is presumably due to the longer DNA linker and missing contacts with DNMT3B3 #2, which binds to the lower nucleosome in the 10-8-10 di-nucleosome (**Fig. 2c-e**). This indicates that di-nucleosome bridging by DNMT3B3 is highly dependent on the DNA linker length, specifically favoring linkers that are between 5-8bp. Linker length dependence of di-nucleosome binding has also been observed for the H3K27 methyltransferase PRC2. However, PRC2 favors long di-nucleosome linkers^40^, which contrasts with the observed preference of DNMT3A2/3B3 for short di-nucleosome linkers. Importantly, the preference of DNMT3A2/3B3 to bind di-nucleosomes with short linkers is not a strict requirement as the hetero-tetramer can still bind di-nucleosomes with long linkers without engaging the second nucleosome (**Fig. 3c**).

### The DNMT3A2/3B3 hetero-tetramer manipulates the di-nucleosome structure

The similar conformations of the 10-5-10, 10-6-10 and 10-8-10 di-nucleosome DNMT3A2/3B3 complexes prompted us to examine how DNMT3A2/3B3 binding may alter the structure of the di-nucleosome substrate itself. We determined Cryo-EM structures of the 10-8-10 and 10-5-10 di-nucleosomes without the DNMT3A2/3B3 hetero-tetramer (**Fig. 4a,b**). Compared to the 10-8-10 di-nucleosome, the 10-5-10 di-nucleosome is more compact, with the lower nucleosome tucked under upper nucleosome, partially occluding its acidic patch (**Fig. 4a,b**). DNMT3A2/3B3 binding to the 10-5-10 di-nucleosome rotates the lower nucleosome along the DNA linker by 23° and away from its tucked in position through a 43° rotation that straightens the DNA linker and moves the acidic patch by 61Å (**Fig. 4c**). These motions allow the DNMT3B3 catalytic-like domain to engage the acidic patch of the lower nucleosome. DNMT3A2/3B3 binding to the 10-8-10 di-nucleosome rotates the lower nucleosome clockwise along the DNA linker by 27° relative to the DNMT-free state (**Fig. 4d**). This rotation moves the acidic patch of the lower nucleosome down by 26Å, enabling DNMT3B3 #2 to interact with the lower nucleosome. These movements may put strain on the di-nucleosome linker because it forces the DNA to adopt a conformation that does not match that of relaxed b-form DNA (**Fig. 3a,b**). This is supported by our cryo-EM structures showing that the linker DNA is better resolved in the structure of the 10-8-10 di-nucleosome alone than in the structure of the 10-8-10 di-nucleosome in complex with DNMT3A2/3B3 (**Fig. S8e,f**), even though the complex structure has a higher overall resolution (**Fig. S5a, S6f**). In the DNMT-free conformation, the DNMT3B3 #2 ADD domain would completely overlap with the lower nucleosome in the 10-5-10 and 10-8-10 di-nucleosome substrates (**Fig. 4c,d**). In addition, a large portion of the DNMT3B3 #2 catalytic-like domain would overlap with the lower nucleosome of the 10-8-10 di-nucleosome in the DNMT-free sate (**Fig. 4d**). The strong clashes with DNMT3B3 #2 in the DNMT-free di-nucleosome structures suggests that the DNMT-free conformation of the 10-5-10 and 10-8-10 di-nucleosomes are incompatible with DNMT3A2/3B3 hetero-tetramer binding. Therefore, the observed conformational changes between the two nucleosomes are required for the DNMT3A2/3B3 hetero-tetramer to bind the di-nucleosome.

**Figure 4:**
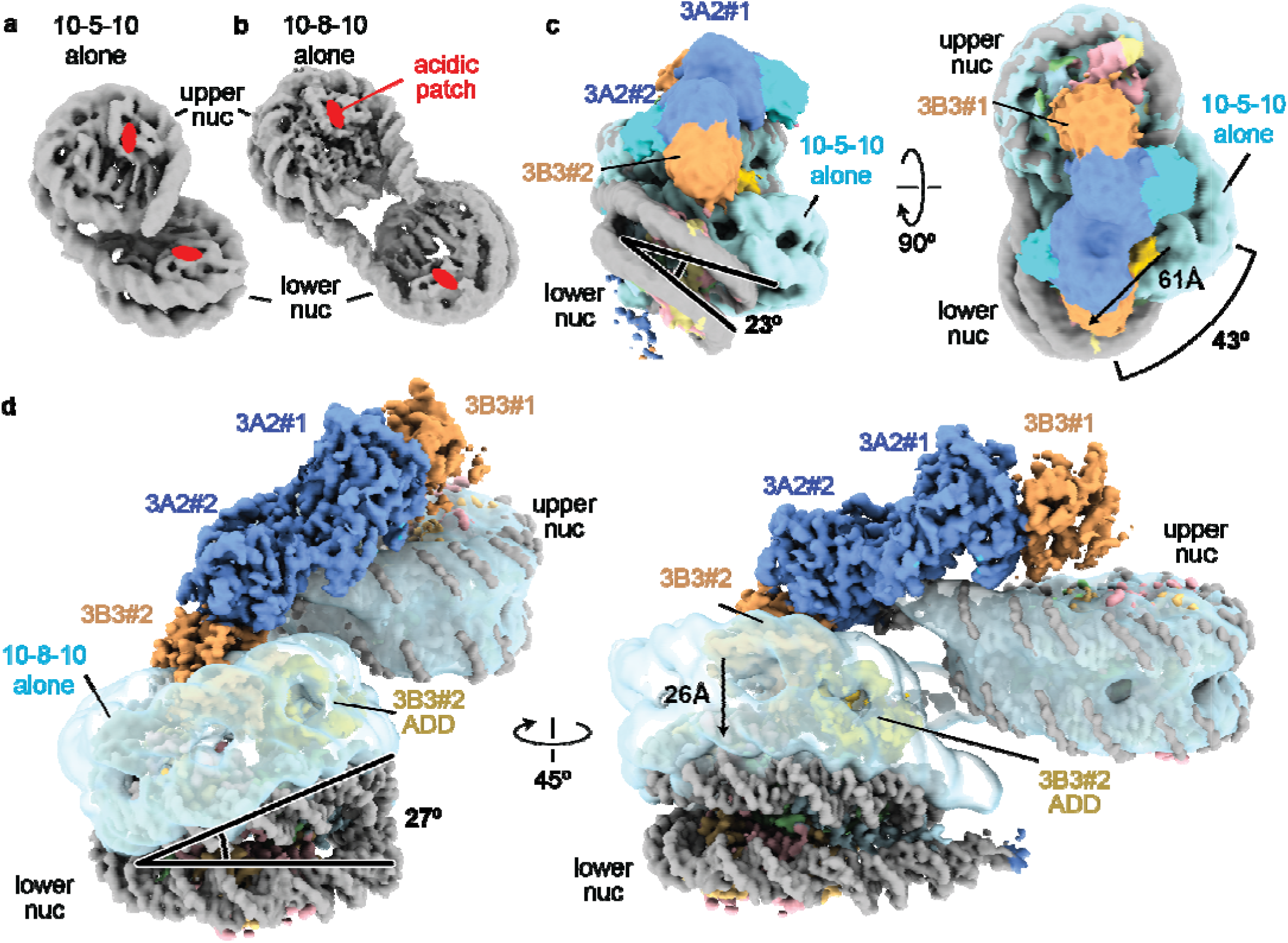
DNMT3A2/3B3 manipulates the conformation of the di-nucleosome. **a-b** Cryo-EM maps for the isolated 10-5-10 (**a**) and 10-8-10 (**b**) di-nucleosomes. The location of the acidic patch in each di-nucleosome is indicated. **c** Consensus cryo-EM map 10-5-10 DNMT3A2/3B3 complex superimposed with the cryo-EM map of the 10-5-10 di-nucleosome alone (light blue). Movements of the lower nucleosome upon DNMT3A2/3B3 tetramer binding are indicated. **d** Composite cryo-EM map of the 10-8-10 DNMT3A2/3B3 complex superimposed with the 10-8-10 di-nucleosome alone depicted as a semitransparent blue surface.

### DNA methylation is regulated by the nucleosome linker length

To systematically dissect the linker length dependence of CpG methylation by DNMT3A2/3B3, we reconstituted a set of di-nucleosome substrates with long overhangs (∼40bp) and linker DNA ranging from 5bp-35bp (**Fig. 5a**). We inserted specific CpG dinucleotides at various positions within the linker DNA and reacted the di-nucleosomes with a 10x excess of DNMT3A2/3B3 in the presence of histone H3 peptide (1-20) and collected endpoints of the reaction to simultaneously assess the effects of linker length and CpG location on DNA methylation using bisulfite sequencing (**Fig. 5b,c**). Our results show that nucleosome spacing and CpG position impose a strong barrier to deposition of linker mCpG by DNMT3A2/3B3. We observed almost no CpG methylation within 5bp, 8bp or 10bp linkers, but as the linker length increased from 15bp to 35bp there was a corresponding increase in linker DNA methylation (**Fig. 5b**). This methylation profile also suggests that DNMT3A2/3B3 primarily deposits hemi-methylation on the nucleosome linker as maximum methylation only approached 50% for both DNA strands. This effect was completely dependent on nucleosomes as there was no correlation between linker length and CpG methylation for identical DNA substrates without histones (**Fig. 5c**). These results indicate that close nucleosome spacing stops DNMT3A2/3B3 from methylating linker DNA and agrees with our Cryo-EM structures of the 10-5-10, 10-6-10 and 10-8-10 di-nucleosome DNMT3A2/3B3 complexes showing that DNMT3A2 does not contact the linker DNA (**Fig. 1f,3a**). Manually generated models of di-nucleosomes separated by 10bp, 15bp or 35bp linkers show that CpG methylation is strongly correlated with steric constraints imposed by the two nucleosomes (**Fig. 5d-g**). Superimposing the mono-nucleosome-bound structure of DNMT3A2/3B3 (PDB: 6PA7) onto the 10bp di-nucleosome model shows that DNMT3B3 #2 would clash with the second nucleosome, potentially explaining the very low linker methylation in this substrate (**Fig. 5d**). This clash is not present the 15bp di-nucleosome model, however linker methylation is still substantially reduced in this substrate relative to free DNA (**Fig. 5b,e**). Reduced linker methylation in the 15bp di-nucleosome is likely due to the overhang DNA that protrudes from the di-nucleosome next to DNMT3A2/3B3 (**Fig. 5e**). Linker methylation would position DNMT3A2/3B3 directly in front of the DNA exit site where the ∼40bp DNA overhangs present in our substrate would directly clash with DNMT3A2 and interfere with linker methylation. Models for the 20bp and 35bp di-nucleosomes superimposed with the mono-nucleosome-bound structure of DNMT3A2/3B3 only result in minor clashes with the DNMT3A2/3B3 hetero-tetramer that could be relieved by small changes in the orientation of DNMT3A2/3B3 (**Fig S9a,b**), agreeing with the higher level of linker methylation in these substrates (**Fig. 5b**). These data indicate that the structure the di-nucleosome imposes strong steric constraints for linker DNA hemi-methylation by DNMT3A2/3B3.

**Figure 5:**
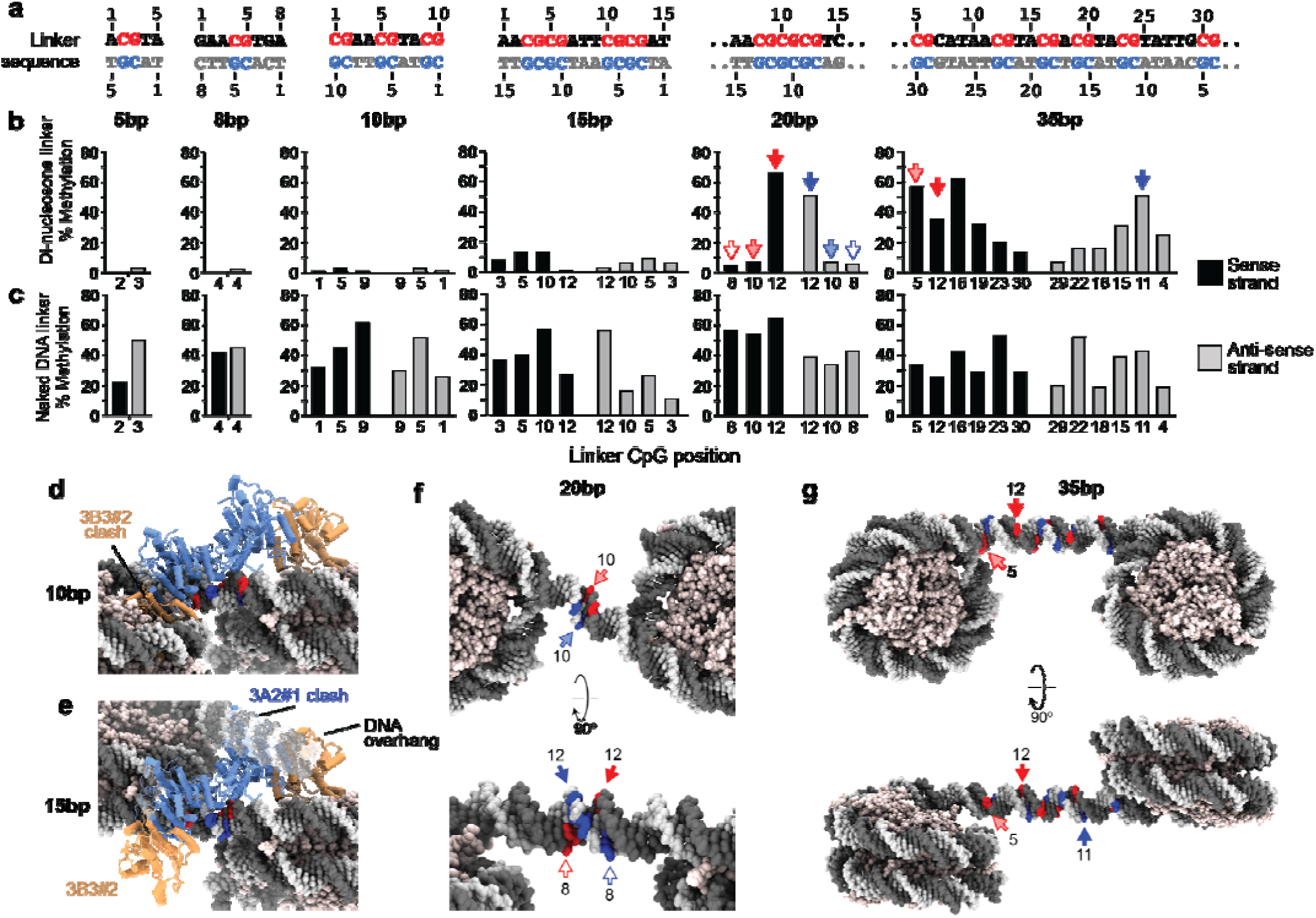
DNA linkers regulate methylation by DNMT3A2/3B3. **a** Sequences of engineered di-nucleosome linkers. Positions of CpG dinucleotides in the sense and antisense strands are numbered based on their location relative to the upstream nucleosome and are colored red and blue, respectively. **b-c** bisulfite sequencing assay showing the percentage of CpG methylation at each CpG position. Specific cytosines are indicated with colored arrows. Top: nucleosome substrates, bottom: naked DNA substrates. **d-g** Manually generated models of the 10bp, 15bp, 20bp and 35bp di-nucleosomes with the positions of target cytosines in the sense and anti-sense strands colored red and blue, respectively. **d-e** Di-nucleosome models superimposed with the structure of mono-nucleosome bound DNMT3A2/3B3 (PDB: 6PA7) to show possible clashes. **f-g** Models of the 20bp and 35bp di-nucleosomes with the positions of specific cytosines indicated using colored arrows.

Deposition of mCpG in 20bp and 35bp linkers shows a pronounced strand-, and position-specific CpG methylation pattern (**Fig. 5b**). Importantly, these effects do not depend on excess H3 peptide used in the reaction, as reactions without added peptide display the same methylation profiles, albeit with lower overall methylation levels (**Fig. S9c**). In the 20bp di-nucleosome substrate, the CpG sites at position 12 in both the sense and anti-sense strands are highly methylated while the other CpG sites have low methylation levels (**Fig. 5b**). Examination of the manually built di-nucleosome model for this substrate shows that the highly methylated C12 on each strand is exposed and perfectly positioned for methylation by the DNMT3A2/3B3 hetero-tetramer (**Fig. 5f**, solid red and blue arrows). However, C10 and C8 are rotated down into the space between the di-nucleosome, where access to the linker is highly restricted, explaining the lower levels of methylation for these CpGs (**Fig. 5b,f**, open and shaded red and blue arrows). The 35bp di-nucleosome exhibits a distinct methylation profile, with high levels of CpG methylation centered at ∼12bp from the upstream nucleosome in the sense strand and at ∼11bp in the anti-sense strand followed by decreased methylation farther away from the upstream nucleosome (**Fig. 5b**). These results agree with biochemical and structural studies suggesting that DNMT3A2/3B3 preferentially methylates mono-nucleosome DNA overhangs ∼10-12bp from the DNA exit site^19,41^. However, we also observed strong methylation of the CpG at position 5 in the sense strand (**Fig. 5b**), which is turned down and away from the expected DNMT3A2/3B3 methylation site (**Fig 5g**, shaded red arrow). This CpG is located at the stiff end of the 601 DNA sequence, which unwraps from the nucleosome with high frequency^42^. Therefore, we attribute the strong methylation at this CpG site to local DNA unwrapping from the nucleosome, which could allow DNMT3A2/3B3 to access CpG #5 for methylation. Together these data show that CpG methylation in the linkers between nucleosomes is tightly regulated by the linker DNA length and by the position of the CpG dinucleotide within the DNA linker itself.

### Short nucleosome linkers are enriched at promotor regions and correlate with DNMT3A2

Because the inter-nucleosome linker length directly controls CpG methylation, we examined how linkers of different lengths are distributed in the genome and are correlated with transcription, DNA methylation and DNMT3A2. We re-analyzed previously published datasets from mouse ES cells reporting on chemically mapped nucleosome linker lengths^32^, ChIP-seq profiles of DNMT3A2, and genome-wide mCpG locations^24^. Mouse ES cells express DNMT3A2 and DNMT3B6^21^, which contains the same catalytic domain deletion as DNMT3B3, suggesting that DNMT3B6 interacts with the nucleosome in a similar way. The linker length distribution at promotor regions of unexpressed genes closely matches the genome wide linker length distribution (**Fig. S10a**). However, at actively transcribed genes the linker length distribution shifts to shorter lengths (**Fig. S10a**). This linker length shortening is concentrated in the 5-20bp length scale and is correlated with transcription, as highly expressed promotors have the shortest nucleosome lengths. Linkers between 5-15bp inhibit DNA methylation (**Fig. 5**), indicating that the enrichment of short linkers at promotor regions may make them naturally resistant to DNA methylation. A much less pronounced linker length shortening is evident for gene bodies, which contain DNA methylation (**Fig. S10a**). This agrees with prior work showing that the nucleosome repeat length of promotors is much shorter than for gene bodies and heterochromatin^33^. Very short nucleosome linkers (0-10bp) are enriched ∼1000bp upstream from the TSS at expressed genes, but not silent genes (**Fig. S10b**). These short linker lengths allow DNMT3A2/3B3 to tightly bind the di-nucleosome by bridging both acidic patches (**Fig. 1**). Interestingly, the enrichment of short linkers at -1000bp co-localizes with a ChIP-seq peaks of DNMT3A2 (**Fig. S10c**). This region also sits at the transition point between the demethylated promotor and methylated intergenic regions (**Fig. S10d**). Together these data suggest that the distribution of nucleosomes with very short linkers at promoter regions correlates with transcription and that DNMT3A2/3B3 may be able to recognize regions enriched in very short DNA linkers that allow di-nucleosome bridging.

### The PWWP domain scans for H3K36me2-3

In all structures of the DNMT3A2/3B3 hetero-tetramer bound to a di-nucleosome, a PWWP domain binds between the ADD domains of DNMT3A2 #1 and DNMT3B3 #2 and is positioned directly at the H3 tail exit site (**Fig. 6a-b**). To confirm that this density is the PWWP domain, we purified the DNMT3A2(476-912)/3B3(Δ234-286) variant that lacks PWWP domains from DNMT3A2 and DNMT3B3 (ΔPWWP). A cryo-EM structure of the DNMT3A2/3B3 ΔPWWP domain hetero-tetramer bound to the 10-8-10 di-nucleosome clearly lacks density spanning the ADD domains of DNMT3A2 #1 and DNMT3B3 #2, confirming that this density corresponds to a PWWP domain from DNMT3A2 or DNMT3B3 (**Fig. S11a**). Because a long, disordered linker connects the PWWP domains of DNMT3A2 and DNMT3B3 to the rest of the hetero-tetramer, it is not possible to determine which DNMT subunit is connected to the PWWP domain. In addition, the EM density for the PWWP domain is weak compared to the neighboring ADDs of DNMT3A2 and DNMT3B3 (**Fig. S11b**) suggesting that the PWWP domain is highly mobile. Indeed, it was not possible to conclusively determine the orientation of the PWWP domain based only on the Cryo-EM density as no secondary structure elements were visible in the map. Therefore, we docked the crystal structure of the DNMT3A PWWP domain (PDB:3LLR) into the Cryo-EM density based on the PWWP domain of LEDGF bound to a nucleosome bearing the H3K36me3 modification (PDB:6S01) followed by ridged body fitting into the density using ChimeraX (See methods, **Fig. 6a-b**). We identified two plausible docking orientations of the PWWP domain with nearly identical fit statistics using this approach (**Fig. S11c-e**). We selected the docking orientation of the PWWP (orientation #1) that placed the H3K36me3 binding site closest to the H3 tail exit point (**Fig. S11c**). The docked model of the PWWP fits well in the low-resolution density and does not clash with the neighboring ADD domains (**Fig. 6a**). The contacts between the PWWP domain and the ADDs of DNMT3A2 and DNMT3B3 are mediated by the helical bundle and beta-sheet of the PWWP domain (**Fig. 6a**). While the precise interacting residues cannot be determined at the resolution of the map, the interfaces between the PWWP and each ADD are small and not predicted by Alphafold2^43,44^ multimer (data not shown). This suggests that each interaction between the PWWP and ADD domains is of low affinity and transient, explaining the weak density for the PWWP domain in our structure (**Fig. S11b**). Indeed, affinity of the DNMT3B PWWP domain for the DNMT3B ADD domain is very low (∼ 52uM)^28^ agreeing with our observation that PWWP domain binds weakly to the ADDs of DNMT3A2 and DNMT3B3. Interestingly, the interfaces between the PWWP domain and each ADD domain in our structure are distinct from the PWWP/ADD interaction in the recently reported Cryo-EM structure of the DNMT3B homo-tetramer (PDB: 8EIH)^28^ (**Fig. S11f-g**). However, the structure of the DNMT3B homo-tetramer was determined in the absence of nucleosomes and the PWWP/ADD interaction was only observed on the terminal ADD domain, not between ADD domains. Therefore, our Cryo-EM structure reports on fundamentally new contacts between the PWWP domain and neighboring ADDs which have important implications for DNMT regulation.

**Figure 6:**
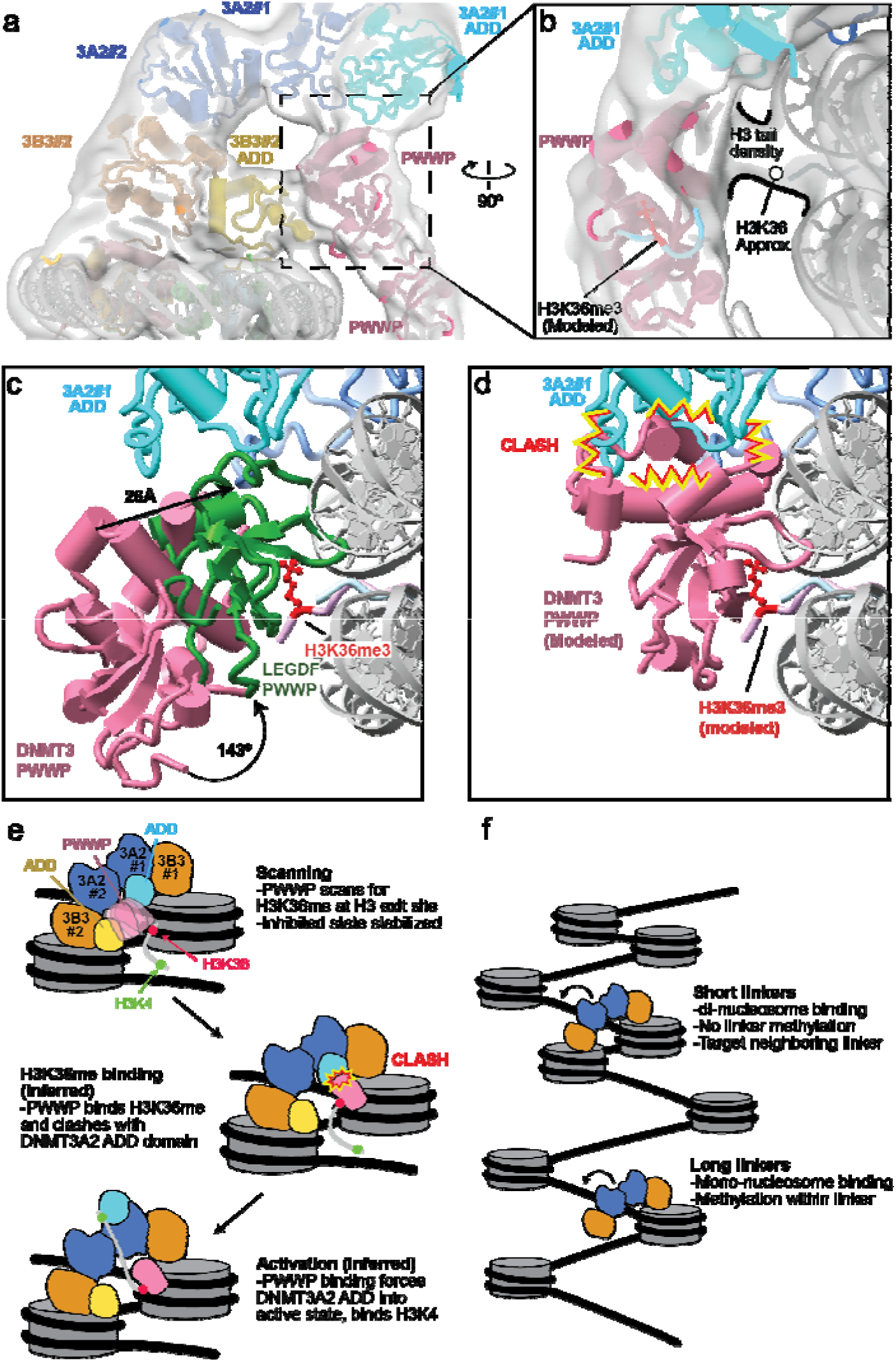
The PWWP domain scans the modification state of H3K36. **a** Model of the DNMT3A2/3B3 complex with the 10-8-10 di-nucleosome docked into the consensus cryo-EM ma of the complex filtered to 10Å. **b** close up view of the PWWP domain to show H3 tail density connecting the PWWP domain to the nucleosome. An H3 peptide from the crystal structure of the DNMT3B PWWP domain bound to H3K36me3 (PDB:5CIU) is modeled to show the location of the H3K36me3 binding site. **c** Structure of the H3K36me3 modified nucleosome bound to the PWWP domain from LEDGF (PDB:6S01) superimposed with the upper nucleosome in the DNMT3A2/3B3 10-8-10 di-nucleosome complex structure. **d** Structure of the DNMT3A2/3B3 10-8-10 di-nucleosome complex with the PWWP domain superimposed with the PWWP from LEGDF (PDB: 6S01, not shown). The helical domain of the PWWP clashes with the ADD domain of DNMT3A2 #1. **e** Model of DNMT activation mediated by PWWP domain recognition of H3K36me2 and ADD domain binding to unmodified H3K4. **f** Model for CpG methylation by DNMT3A2/3B3 in the context of chromatin.

The position of the PWWP domain suggests that it is poised for recognition of the H3K36me2 modification, which is located directly at the H3 tail exit site (**Fig. 6b**). Low resolution density for the unmodified H3 tail connecting to the PWWP domain is visible in the structure (**Fig. 6b**). However, the approximate location of H3K36 is far away from the H3K36me3 binding site on the PWWP domain (**Fig. 6b, S11c**), indicating that the PWWP domain is not bound to H3K36. Furthermore, the position of the DNMT3 PWWP domain differs from the position of the related LEDGF PWWP domain when bound to a nucleosome containing the H3K36me3 modification (**Fig. 6c**)^45^. Therefore, the DNMT3 PWWP domain must reposition itself to recognize the H3K36me2 modification. The position of the LEGDF PWWP indicates that the DNMT3 PWWP domain would need to dissociate from the DNMT3A2 #1 and DNMT3B3 #2 ADD domains and move toward the nucleosome by ∼26Å and rotate by ∼143° to bind to the H3K36me2-3 modification (**Fig. 6c**). The low resolution and high mobility of the PWWP domain indicates that it may scan the modification state of H3K36 by periodically breaking its interactions with the ADDs of DNMT3A2 and DNMT3B3. H3K36 is unmodified in our structure, and we could not identify any class of particles where the PWWP domain engages H3K36 in our dataset. This indicates that any excursions of the PWWP domain to scan the modification state of H3K36 are short lived and likely require H3K36me2-3 for stabilization. Importantly, this scanning conformation of the PWWP domain does not depend on di-nucleosome bridging, because weak PWWP density is also evident in the structure of DNMT3A2/3B3 bound to a mono-nucleosome^19^ (**Fig. S11h**). Together, our structure shows that the DNMT3A2/3B3 hetero-tetramer optimally positions a PWWP domain to read the modification state of H3K36, allowing the DNMT to identify correctly modified nucleosome substrates for DNA methylation.

### H3K36 recognition by the PWWP clashes with the inhibited ADD conformation

DNMT3A activity is allosterically stimulated by the H3 tail containing unmodified H3K4, which binds the ADD domain and changes its conformation to a “flipped-up” state^22^ (**Fig. S7b-c**). Adding excess H3 (1-20) peptide to the DNMT3A2/3B3 10-8-10 di-nucleosome complex did not affect the conformation of the DNMT3A2 ADD domains (**Fig. S11i**), suggesting that nucleosome binding stabilizes the auto-inhibited conformation of the DNMT3A2 ADD domains. Stabilization of the DNMT3A2 ADD domain in the auto-inhibited conformation may be enhanced by the PWWP domain, which bridges the ADD domains of DNMT3A2 #1 and DNMT3B3 #2 in the inactive state (**Fig. 6a**). Furthermore, PWWP mutations and deletions increase DNMT3A methylation activity, suggesting that the PWWP domain inhibits DNMT3A activity^28,46^. DNMT3A and DNMT3B are also activated upon recognition of H3K36me2 by the PWWP domain^29^, indicating that H3K36me2 binding to the PWWP domain can release its inhibitory effect. Our structure shows that recognition of H3K36me2-3 requires large-scale movement of the PWWP domain that would break contacts with the ADDs of DNMT3A2 and DNMT3B3, potentially destabilizing the inhibited state conformation (**Fig. 6c-d**). In addition, superimposing the DNMT3B PWWP domain onto the PWWP domain of LEDGF^45^ shows that PWWP binding to H3K36me2-3 would clash with the DNMT3A2 #1 ADD domain in the autoinhibited state (**Fig. 6d**). Therefore, movement of the DNMT3A2 #1 ADD domain away from the autoinhibited state would be required to accommodate PWWP binding to H3K36me2-3. Together, these data indicate that nucleosome binding stabilizes the ADD domains of DNMT3A2 in an inhibited conformation and that recognition of H3K36 methylation by the PWWP domain may destabilize the inhibited state of DNMT3A2 to activate the enzyme for CpG methylation.

## Discussion

Our study shows that the DNMT3A2/3B3 hetero-tetramer preferentially binds to di-nucleosomes separated by very short DNA linkers (<10bp) by engaging the acidic patches of both nucleosomes with arginine anchors within DNMT3B3. In the di-nucleosome bound state, DNMT3A2 does not bind the linker DNA and instead binds to the DNA overhangs, which would correspond to neighboring DNA linkers in the context of chromatin. Therefore, closely spaced di-nucleosomes with short linkers may serve as high affinity DNMT3A2/3B3 recruitment signals to direct DNA methylation to neighboring linkers. Recent studies have shown that the N-terminal region of DNMT3A1 (which is missing in DNMT3A2) binds to nucleosomes modified with H2AK119Ub^14^. While there are no structures of hetero-tetramers containing DNMT3A1 bound to the nucleosome, structures of the N-terminal region of DNMT3A1 show that it forms extensive contacts with the nucleosome acidic patch^29,47,48^. We expect that the extensive contacts between the DNMT3A1 N-terminal region and the acidic patch would compete with DNMT3B3 and its absence in DNMT3A2 probably allows DNMT3B3 to bind the nucleosome as observed in our structures. Therefore, the role of short linker di-nucleosomes as a DNMT recruitment signals may depend on which DNMT3A isoform is expressed. DNMT3A1 is the primary catalytic subunit expressed in differentiated tissues^20^. This indicates that the di-nucleosome bridging observed our structure may not predominate in somatic cells where DNMT3A1 is more highly expressed than DNMT3A2^19,20^. However, in ES cells and in various cancers DNMT3A2 is highly expressed^19^. Therefore, the interaction between the DNMT3A2/3B3 hetero-tetramer and short linker di-nucleosomes captured in our structure may be particularly important in the context of cancer and stem cell development.

The structures of the 10-5-10 and 10-8-10 di-nucleosomes alone and in complex with DNMT3A2/3B3 show that hetero-tetramer binding dramatically alters the conformation of the di-nucleosome. In each case, DNMT3A2/3B3 binding forces the di-nucleosome toward a single conformation that places the acidic patch of the lower nucleosome in direct contact with DNMT3B3#2. Cryo-EM structures of the 10-5-10 and 10-8-10 di-nucleosomes in the DNMT-free state occupied a single defined conformation, suggesting that the natural fluctuations in these di-nucleosomes are very rare or not significant enough to access the DNMT3A2/3B3 bound conformations. Therefore, we expect that the DNMT may actively push the di-nucleosome away from its ground state conformation to a single DNMT bound state and pay the energetic cost of deforming the di-nucleosome linker DNA using binding energy from interactions with the acidic patch and the DNA overhangs. This may enable the DNMT3A2/3B3 hetero-tetramer to bind di-nucleosomes within a range of linker lengths so that it can precisely position the lower nucleosome close to the DNMT3B3 arginine anchors. This contrasts with other di-nucleosome binding proteins like PRC2^49^, which adapts to changes in linker length by accommodating different nucleosome conformations and does not force the di-nucleosome to adopt a single bound state conformation. Therefore, DNMT3A2/3B3 may use a unique mechanism to accommodate changes in the di-nucleosome linker length by directly manipulating the conformation of the di-nucleosome toward a single DNMT bound state.

Di-nucleosome linker DNA methylation by DNMT3A2/3B3 revealed a striking connection between DNA linker length, CpG positioning and cytosine methylation frequency (**Fig. 5**). Surprisingly, linkers shorter than 15bp were weakly methylated, even under the highly favorable conditions employed in our methylation assay. This suggests that DNA linker length directly regulates the level of linker DNA methylation, with closely spaced nucleosomes strongly inhibiting linker DNA methylation. In mice and humans, highly expressed genes are enriched with shorter linkers the 5-20bp range^32^ and this linker length shortening is concentrated near promotors of active genes (**Fig. S10a**)^33^, which contain low levels of DNA methylation in normal cells. In contrast, DNA linkers longer than 35bp are concentrated in gene bodies and within heterochromatin^32,33^, which contain high levels of DNA methylation. The enrichment of shorter DNA linkers near promoters of active genes indicates that these regions may have natural resistance to DNA methylation *in vivo*, while longer linkers within heterochromatin and gene bodies may permit unhindered DNA methylation. Interestingly, DNA linkers in the 0-10bp length range, which allows di-nucleosome bridging by DNMT3A2/3B3 (**Fig. 1**), are concentrated ∼1000bp upstream from transcription start sites at the transition point between highly methylated intergenic regions and poorly methylated promotors. Linkers in this length range also colocalize with DNMT3A2, indicating that the specific positioning of very short linkers may be a mechanism used by the cell to recruit DNMT3A2 to genomic loci. While our structural studies show how methylation is regulated within nucleosome linkers, it is still not understood how the DNA that wraps the nucleosome is methylated. One possible explanation is that methylation of nucleosomal takes place concurrently with nuclear processes that disrupt nucleosome positioning, such as replication, transcription, and nucleosome remodeling. Importantly, nucleosome linker lengths are regulated by the action of chromatin remodeling complexes, which are critical to enable DNA methylation^30^. Therefore, regulation of the inter-nucleosome DNA linker length could be a mechanism used by the cell to direct DNA methylation away from active gene promotors and toward heterochromatin and gene bodies.

We now propose a model of DNMT3A2/3B3 activation and methylation where the PWWP domain has dual roles in regulating DNMT3A2 activity, first by stabilizing the inhibited conformation of the DNMT3A2 ADD domain in the absence of H3K36me2-3, followed by destabilizing the inhibited conformation upon recognition of H3K36me2-3 (**Fig 6e**). Upon initial nucleosome engagement, the DNMT3A2/3B3 hetero-tetramer adopts an inhibited conformation where the DNMT3A2 ADDs are in the “flipped-down” state and stabilized by the PWWP domain which bridges the ADD domains of DNMT3B3 and DNMT3A2. The PWWP domain is positioned at the H3 exit site to scan for H3K36me2-3 modifications (**Fig. 6e**, scanning). Recognition of H3K36me2-3 by the PWWP domain destabilizes the inhibited state by breaking contacts between the PWWP domain and each ADD domain, and by inducing a clash between the PWWP domain and the DNMT3A2 ADD domain (**Fig. 6e**, H3K36me binding). This clash between the PWWP domain and the DNMT2A2 #1 ADD domain may force the ADD domain into the “flipped-up” conformation and activate the enzyme if H3K4 is also unmodified (**Fig. 6e**, activation). CpG methylation then occurs in the linker DNA between nucleosomes, however the level of linker methylation depends on the corresponding linker length and position of the CpG dinucleotide (**Fig. 6f**). Short linkers (<15bp) between nucleosomes are resistant to DNA methylation and very short linkers (5-8bp) enable di-nucleosome bridging by the DNMT3A2/3B3 hetero-tetramer and completely inhibit linker methylation. In contrast, long linkers enable efficient CpG methylation due to the long distance between nucleosomes (**Fig. 6f**). Together, our structural and biochemical studies provide a mechanistic framework to understand the regulation of DNMT3A2/3B3 hetero-tetramers in chromatin and explain the dual requirement for unmodified H3K4 and H3K36me2-3 for DNMT3 activation and CpG methylation.

## Author Contributions

X.X., P.A.J. and E.J.W. conceived of the study. X.X., M.L. and M.L.D. performed experiments. X.X. processed Cryo-EM data and X.E.Z. refined the atomic model. E.J.W. wrote the manuscript with assistance from X.X., M.L. and P.A.J.. P.A.J and E.J.W. supervised the study.

## Acknowledgements

This work was supported by the National Institutes of General Medical Sciences under award R35GM147261 (E.J.W.) and by the National Cancer Institute under awards R35CA209859 (P.A.J.) and R50CA243878 (M.L.).

## Methods

### DNMT3A2/DNMT3B protein preparation

The human proteins DNMT3A2 and DNMT3B3 were expressed and purified from SF9 insect cells as described previously^19^. Briefly, P1 viruses containing DNMT3A2 and 8xHis-GFP-TEV-DNMT3B3 were used to co-infect SF9 cells and express proteins. The proteins were purified by using a nickel-NTA affinity column. After the removal of the 8xHis-GFP tag by addition of TEV protease, the proteins were further purified using a Superdex 200 size exclusion column. The peak containing the DNMT3A2/3B3 hetero-tetramer was collected, concentrated, and flash-frozen in liquid nitrogen and stored at -80°C.

### Histone octamer preparation and Di-nucleosome reconstitution

Soluble *X. laevis* histone octamers containing 6xHis-SUMO-H2A, H2B, 6xHis-SUMO-H3, and H4 were purified as described previously^19^. Briefly, the histone proteins were co-expressed in E. coli and purified by nickel-NTA affinity chromatography. The 6xHis-SUMO tag was removed by ULP1 protease and the histone proteins were further purified using a Hitrap SP cation exchange chromatography column. The proteins were loaded and washed on the column using buffer A (20mM Tris PH8.0, 500 mM NaCl,10% glycerol, 5mM B-Me) and eluted with a 0% - 100% linear gradient of buffer B (20mM Tris PH8.0, 2M NaCl,10% glycerol, 5mM β-mercaptoethanol) over 6 column volumes (CV). The histones were further purified a Superdex 200 size exclusion column (Cytiva) and peaks containing the desired octamer were pooled and concentrated. The purified octamer was flash-frozen in liquid nitrogen and stored at -80°C.

### Preparation of Di-nucleosome DNA fragments

The di-nucleosomal DNA fragments contain two copies of the Widom 601 nucleosome positioning sequence^50^ separated by a variable linker sequence and is flanked by defined overhang DNAs and EcoRV cut sites to excise the DNA from the vector. All DNAs were synthesized and cloned into pBluescript II KS (-) using Genscript custom gene synthesis service. OmniMAX 2 T1R cells (Thermofisher) were transformed with the plasmid containing the di-nucleosome DNA and plated on ampicillin plates. 30 ml of LB medium containing ampicillin (100 μg/ml) was inoculated with single colony and cultured overnight at 37 °C while shaking at 220 rpm. For the DNA growth, 10 ml of preculture was transferred into to each 4L flask containing 1000 ml of terrific broth (TB) and ampicillin at 100 μg/ml. The cells were incubated under vigorous shaking for 18 h at 37°C and harvested by centrifugation in 500-ml centrifuge bottles. Plasmid was extracted as described previously^51^. After RNase A digestion, the plasmid solution was centrifuged at 17,500 rpm for 30 min at 4 °C. The supernatant containing plasmid DNA was filtered and loaded onto a 5-ml HiTrap Q column pre-equilibrated with buffer A (10 mM Tris, pH 8.0, 1 mM Na-EDTA) and washed with 10 CVs of 30% buffer B (10 mM Tris, pH 8.0, 1 mM Na-EDTA). The plasmid was then eluted with a linear NaCl gradient from 300 mM to 1M over 6 CVs. Each fraction was analyzed by 1% (w/v) TAE agarose gel electrophoresis. Fractions with plasmid-DNA were pooled and precipitated by the addition of 1/10 volume of 3M sodium acetate PH 5.2 and 2.5 volumes of 100% ice-cold ethanol. The plasmid DNA pellet was air dried for 30 min at room temperature and dissolved in 10 ml TE 10/0.1 buffer (10 mM Tris-HCl, pH 8.0, 0.1 mM EDTA).

The di-nucleosome DNA insert was excised by addition of EcoRV at 37°C for 20h and complete digestion was confirmed by 6% PAGE analysis. The excised insert was separated from the linearized vector by PEG precipitation as described^51^. 7.5% PEG 6000 effectively precipitated vector DNA, with most of the insertion fragments presented in the supernatant. After precipitation in 3M sodium acetate PH 5.2 and 2.5 volumes of 100% ice-cold ethanol the di-nucleosome DNA pellet was air-dried for 30 min at room temperature and dissolved in 10 ml TE 10/50 buffer (10 mM Tris-HCl, pH 8.0, 50 mM EDTA). The insertion fragment was further purified using a HiTrap Q column with the same elution procedure as the plasmid DNA purification. Fractions containing the di-nucleosome DNA were precipitated and the DNA pellet was air-dried for 10 minutes, dissolved with TE 10/0.1 buffer, and then stored at -80°C. The sequences of the prepared template DNA fragments are listed below. The linker and overhang DNA sequences are indicated in bold with underline. 10NxN10 sequences were used for structural studies and the 40NxN38 were used for DNA methylation assays and bisulfite sequencing.

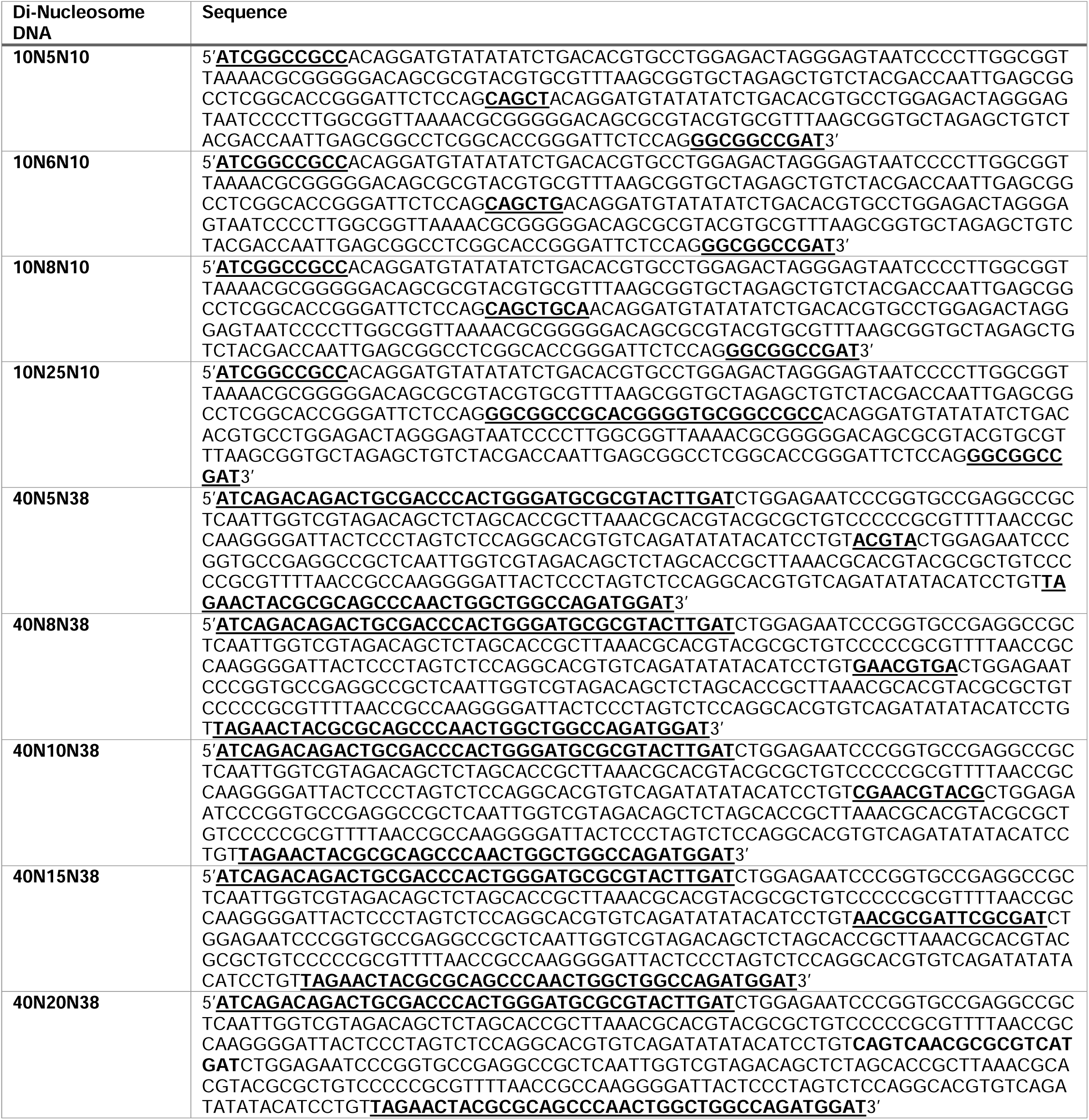
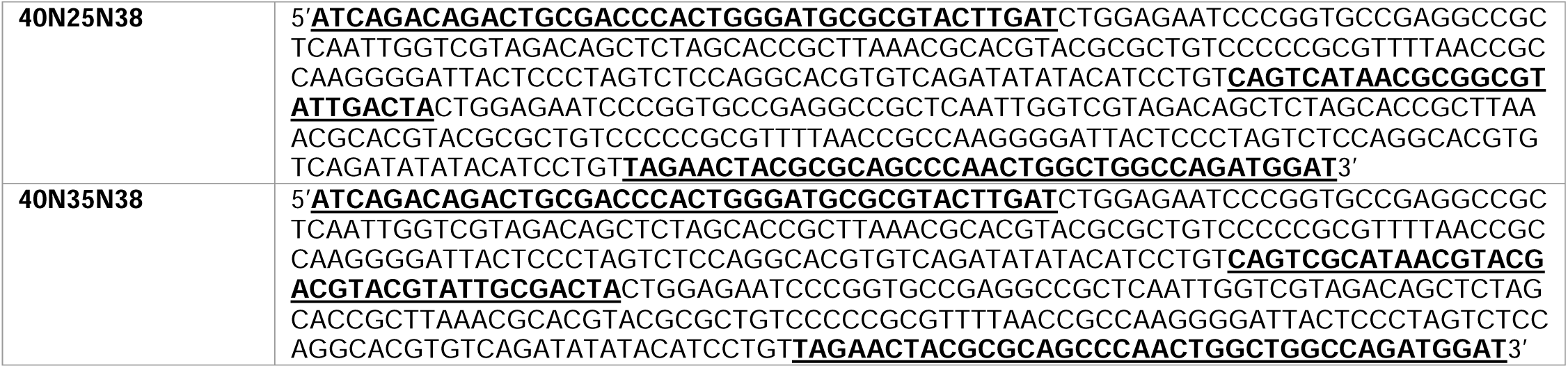

### Di-nucleosome reconstitution

Di-nucleosomes were reconstituted as described previously^51^. Briefly, to reconstitute di-nucleosomes, histone octamers and purified di-nucleosome DNA were mixed at a 2.1:1 molar ratio under high salt. Then, the mixture was dialyzed in high-salt buffer for 4 hours followed by gradient dialysis into low salt buffer over 22 hours.

### Electrophoretic mobility shift assays

125 nM Di-nucleosomes were mixed with the DNMT3A2/3B3 hetero-tetramer (final concentrations ranging from 0 to 500 nM) in EMSA binding buffer(20 mM HEPES pH 7.8, 50 mM KCl, 1 mM MgCl2, 1 mM DTT, 0.1 μM S-adenosyl homocysteine (SAH), 0.1 mg/ml bovine serum albumin, and 5% glycerol). The final volume of the binding reaction was 30 μl. The binding reactions were incubated on ice for 30 minutes and then were examined on a 6% polyacrylamide native gel running in 0.5 x tris-borate EDTA on ice. The gels were stained with CYBR Safe DNA stain (thermofisher) for 15 minutes before being imaged.

### DNA methylation assay and bisulfite sequencing

To assess the activity of the DNMT3A2/3B3 complex on di-nucleosomes with variable linker lengths, 10 pmol of each di-nucleosome (100nM) or its corresponding naked DNA was incubated with 100 pmol of purified DNMT3A2/3B3 hetero-tetramer (1μM) in 100 μl binding buffer (20 mM HEPES pH7.8, 50 mM NaCl, 1% glycerol, 1 mM DTT, 0.1 mg/ml BSA, 100 μM SAM, and 1 μM histone H3 peptides (residues 1–20)) at 37 °C for 4 hours. To remove free DNA from the reaction with the di-nucleosomes, one unit of AluI (New England Biolabs) was added to the reaction and incubated at 37 °C for 15 minutes. The 601 DNA sequence contains a single AluI site that is blocked by histone binding, allowing for specific cleavage of free DNA not bound to the nucleosome. All reactions were then terminated by heating at 80°C for 20 minutes. The DNA was then immobilized on Agencourt AMPure XP Beads (Beckman Coulter), washed twice with 80% ethanol, eluted with water, and subjected to bisulfite conversion using the EZ DNA Methylation Kit (Zymo Research) following the manufacturer’s instructions. Bisulfite-converted DNA was then amplified by PCR using the ZymoTaq DNA Polymerase (Zymo Research) with the following primers: forward, 5′-GATAGATAGTTGTTGAATTAATGGGATT-3′, and reverse, 5′-CTTCATCTTACCAACCAATTAAACAA-3′. PCR was carried out with the following conditions: 95 °C for 3 minutes, followed by 15 cycles of 95 °C for 10 seconds, 55 °C for 30 seconds, and 72 °C for 1 minute, with a final incubation at 72 °C for 5 minutes. PCR products were then sent out for next-generation sequencing (Genewiz) using the Amplicon-EZ service. The sequencing data were analyzed using BISCUIT software (https://huishenlab.github.io/biscuit/).

### Cryo-EM sample preparation for Di-nucleosome & DNMT3A2/3B3 complexes

To prepare the cryo-EM sample for DNMT3A2/3B3-dinucleosome complexes, di-nucleosomes were mixed with DNMT3A2/3B3 at a ratio of 1:2.1. The final concentration of DNMT3A2/3B3 hetero-tetramer was typically between 1.6-2.0 μM. The mixture was then incubated and dialyzed into binding buffer (25 mM HEPES, pH 7.8, 50 mM NaCl, 1 mM DTT, 100 μM SAH) for 2-4 hours. The complex was subjected to GraFix^52,53^ in GF buffer (25 mM HEPES, pH 7.8, 30 mM NaCl, 1 mM DTT, 100 μM SAH) with a glycerol gradient of 10-30% and glutaraldehyde gradient of 0-0.15%. The gradient was fractionated from bottom to top with 500 ul for each fraction. All fractions were checked using a 6% polyacrylamide native gel running in 0.5 x tris-borate EDTA on ice. Fractions containing the desired DNMT3A2/3B3 & di-nucleosome complex were pooled and dialyzed into sample buffer (25 mM HEPES, pH 7.8, 50 mM NaCl, 1 mM DTT) overnight, concentrated and stored on ice. Cryo-EM grids were prepared by incubating 3 μl of the sample for 1 second on glow-discharged 1.2/1.3 or 2/1 Quantifoil holey carbon grids (Quantifoil Micro Tools, Jena) in the humidity chamber of a Mark IV vitrobot (FEI) maintained at 6°C and 100% humidity. The sample was then blotted for 2.5 seconds before being plunged into liquid ethane.

### Data acquisition and image processing of 10n8n10 Di-nucleosome & DNMT3A2/3B3

Cryo-EM movies for 10n8n10 di-nucleosome & DNMT3A2/3B3 complex were collected at 300-kV using a Titan Krios electron microscope equipped with a K3 direct electron detector (Gatan) in super-resolution mode. The defocus range was -1.1 to -1.5 μm, with a nominal magnification of ×105,000, resulting in a calibrated super-resolution pixel size of 0.414 Å. 50 frames were recorded for each movie with a total dose of 61 e/A^2^. 19,437 micrographs were collected and processed. All data sets were processed with CryoSPARC^54^.

CryoSPARC Live installed in a local workstation was used to monitor data collection and preprocess movies during collection. Image preprocessing included patch motion correction (crop factor of 2, pixel size 0.828) and contrast transfer function (CTF) correction. Blob-picking picked particles (200 - 350 Å diameter) were extracted with a binning factor of 4, subjected to 2D classification and good classes were used as templates for further picking. To remove poor-quality images, we set thresholds of CTF fit resolution as 15 Å and ice thickness as 1.16. A total of 18,565 raw micrographs were accepted using these thresholds. Template picking was performed with the accepted images with a reference generated from blob-picking particles. About 3.19 million particles were extracted with a binning factor of 4, and subjected to 2D image classification. Good classes were selected and pooled to result in 2.58 million particles. Multiple round of 3D Ab-initio and heterogeneous refinement ultimately yielded a cleaned particle stack 408,670 unbinned particles. A further heterogeneous refinement job with 2 outputs was performed with 408,670 particles. One of the classes (211,221 particles) showed slightly better density for the PWWP domain and was used as a consensus map for figure preparation.

For particle subtraction and focused refinement, all 408,670 particles were pooled and refined to a resolution of 3.75 Å using the non-uniform refinement job in cryoSPARC. Particle subtraction was performed using a mask around regions of the map that were not included in the focused refinement. After particle subtraction, four separate focused refinements were performed using masks encompassing each DNMT3A2/3B3 hetero-tetramer and each nucleosome. The local resolution of all reconstructions were assessed using CryoSPARC local resolution estimation.

The composite map of the 10-8-10 di-nucleosome DNMT3A2/3B3 structure was generated by combining the unsharpened, and DeepEMhancer^55^ sharpened maps together using the Vop Maximum command in chimera. First, focused maps from each nucleosome were individually combined with a scale factor of 1 for the unsharpened maps and a scale factor of 0.85 for the DeepEMhancer map. Then the focused maps from each DNMT tetramer were individually combined with a scale factor of 1 for the unsharpened maps and a scale factor of 0.9 for the DeepEMhancer maps. The Combined maps for the nucleosomes and the DNMT tetramers were then combined using Vop Maximum with a scale factor of 1.1 for each nucleosome and 0.8 for each DNMT tetramer.

### Data acquisition and data processing of Di-nucleosomes with different linker lengths

Movies of the 10-6-10 Di-nucleosome DNMT3A2/3B3 complex and the 10-8-10 Di-nucleosome (H2BK120R) DNMT3A2/3B3 complex were collected at 300 keV using a Titan Krios electron microscope as described for the 10-8-10 structure. All other samples were collected at 200 keV using a Talos Arctica electron microscope equipped with a K2 direct electron detector with a calibrated super resolution pixel size of 0.46Å.

The Di-nucleosome & DNMT3A2/3B3 complexes with different DNA linkers were processed with CryoSPARC. First, blob-picking was used to acquire the initial particle template. Then, template-based automated particle picking was used to select more particles. Following 2D classification, the desired particles were chosen and subjected to 3D ab initio modeling. The particle set was cleaned using several rounds of heterogeneous refinement. Further refinements, such as homogeneous refinement, CTF refinement, and non-uniform refinement, were utilized to obtain the final maps.

### Model building for DNMT3A2/3B3 di-nucleosome complexes

#### DNMT3A2/3B3 10-8-10 di-nucleosome complex composite map

The model was built into the composite map that was generated from four focused maps. The starting models for the individual subunits were taken from PDB entries 6PA7^19^ (DNMT3A2/3B3, histones, and 601 nucleosome DNA sequence), 4U7P^22^ (DNMT3A ADD) and 3LLR^56^ (PWWP, citation). These models were docked into the EM maps using UCSF Chimera^57^ and rigid body-refined using PHINEX^58^, followed by manually adjustment and rebuilding in COOT^59^. The DNA linker region was manually generated and fitted into the density in COOT. The DNMT3B3 catalytic-like domain and the arginine finger peptides that interact with the nucleosome acidic patch were initially adopted from the model 6PA7 and were refined against the composite map. However, the regions of the connecting loop between the CLD core domain and the arginine finger peptide was truncated due to the disordered density. The whole model was refined against the composite map using phenix.real_space_refine in PHENIX.

#### DNMT3A2/3B3 10-8-10 di-nucleosome complex consensus map

The DNMT3A2/3B3 10-8-10 di-nucleosome complex model generated from the composite map was docked into the consensus map. The PWWP domains and the ADD domains of DNMT3A2 #2, which were not visible in the composite map, were docked in to the EM density, corrected with Coot and refined against the map using Phenix.

#### Focused maps for nucleosome 1, nucleosome 2, DNMT tetramer 1 and DNMT tetramer 2

Each portion of the DNMT3A2/3B3 10-8-10 di-nucleosome complex composite model was used to dock into nucleosome 1, nucleosome 2, DNMT tetramer 1 and DNMT tetramer 2 focus maps, respectively, and refined in PHENIX.

#### 10-8-10 and 10-5-10 di-nucleosomes alone

The each nucleosome from the DNMT3A2/3B3 10-8-10 di-nucleosome complex model was docked into the EM map and the linker region was manually built in COOT. The final models was refined against the EM density in PHENIX.

#### DNMT3A2/3B3 10-8-10 di-nucleosome complexes with H3 peptide, H2B K120R, or PWWP deletion

The DNMT3A2/3B3 10-8-10 di-nucleosome complex model from the composite map was docked into the respective density maps. Models were refined with PHENIX against the respective EM density maps.

#### The DNMT3A2/3B3 di-nucleosome complexes with 10-5-10, 10-6-10, or 10-25-10 linkers

Each nucleosome each DNMT from the10N8N10 di-nucleosome complex model docked into respective EM maps. The DNA linker regions were manually re-built to fit into each density map in COOT. The models were manually re-adjusted and refined in PHENIX.

All the EM map statistics and refinements were listed in Table 1. All figures were prepared in UCSF ChimeraX^57^.

## References

1 Keshet, I., Yisraeli, J. & Cedar, H. Effect of regional DNA methylation on gene expression. Proc Natl Acad Sci U S A 82, 2560–2564, doi:10.1073/pnas.82.9.2560 (1985).

2 Schnedl, W., Erlanger, B. F. & Miller, O. J. 5-methylcytosine in heterochromatic regions of chromosomes in Bovidae. Hum Genet 31, 21–26, doi:10.1007/BF00270395 (1976).

3 Scarbrough, K., Hattman, S. & Nur, U. Relationship of DNA methylation level to the presence of heterochromatin in mealybugs. Mol Cell Biol 4, 599–603, doi:10.1128/mcb.4.4.599-603.1984 (1984).

4 Li, E., Bestor, T. H. & Jaenisch, R. Targeted mutation of the DNA methyltransferase gene results in embryonic lethality. Cell 69, 915–926, doi:10.1016/0092-8674(92)90611-f (1992).

5 Okano, M., Bell, D. W., Haber, D. A. & Li, E. DNA methyltransferases Dnmt3a and Dnmt3b are essential for de novo methylation and mammalian development. Cell 99, 247–257, doi:10.1016/s0092-8674(00)81656-6 (1999).

6 Ehrlich, M. & Ehrlich, K. C. Effect of DNA methylation on the binding of vertebrate and plant proteins to DNA. EXS 64, 145–168, doi:10.1007/978-3-0348-9118-9_7 (1993).

7 Blattler, A. & Farnham, P. J. Cross-talk between site-specific transcription factors and DNA methylation states. J Biol Chem 288, 34287–34294, doi:10.1074/jbc.R113.512517 (2013).

8 Yin, Y. et al. Impact of cytosine methylation on DNA binding specificities of human transcription factors. Science 356, doi:10.1126/science.aaj2239 (2017).

9 You, J. S. & Jones, P. A. Cancer genetics and epigenetics: two sides of the same coin? Cancer Cell 22, 9–20, doi:10.1016/j.ccr.2012.06.008 (2012).

10 Hermann, A., Goyal, R. & Jeltsch, A. The Dnmt1 DNA-(cytosine-C5)-methyltransferase methylates DNA processively with high preference for hemimethylated target sites. J Biol Chem 279, 48350–48359, doi:10.1074/jbc.M403427200 (2004).

11 Robertson, K. D. et al. The human DNA methyltransferases (DNMTs) 1, 3a and 3b: coordinate mRNA expression in normal tissues and overexpression in tumors. Nucleic Acids Res 27, 2291–2298, doi:10.1093/nar/27.11.2291 (1999).

12 Kato, Y. et al. Role of the Dnmt3 family in de novo methylation of imprinted and repetitive sequences during male germ cell development in the mouse. Hum Mol Genet 16, 2272–2280, doi:10.1093/hmg/ddm179 (2007).

13 Xie, S. et al. Cloning, expression and chromosome locations of the human DNMT3 gene family. Gene 236, 87–95, doi:10.1016/s0378-1119(99)00252-8 (1999).

14 Gu, T. et al. The disordered N-terminal domain of DNMT3A recognizes H2AK119ub and is required for postnatal development. Nat Genet 54, 625–636, doi:10.1038/s41588-022-01063-6 (2022).

15 Hata, K., Okano, M., Lei, H. & Li, E. Dnmt3L cooperates with the Dnmt3 family of de novo DNA methyltransferases to establish maternal imprints in mice. Development 129, 1983–1993 (2002).

16 Weisenberger, D. J. et al. Identification and characterization of alternatively spliced variants of DNA methyltransferase 3a in mammalian cells. Gene 298, 91–99, doi:10.1016/s0378-1119(02)00976-9 (2002).

17 Saito, Y. et al. Overexpression of a splice variant of DNA methyltransferase 3b, DNMT3b4, associated with DNA hypomethylation on pericentromeric satellite regions during human hepatocarcinogenesis. Proc Natl Acad Sci U S A 99, 10060–10065, doi:10.1073/pnas.152121799 (2002).

18 Jia, D., Jurkowska, R. Z., Zhang, X., Jeltsch, A. & Cheng, X. Structure of Dnmt3a bound to Dnmt3L suggests a model for de novo DNA methylation. Nature 449, 248–251, doi:10.1038/nature06146 (2007).

19 Xu, T. H. et al. Structure of nucleosome-bound DNA methyltransferases DNMT3A and DNMT3B. Nature, doi:10.1038/s41586-020-2747-1 (2020).

20 Duymich, C. E., Charlet, J., Yang, X., Jones, P. A. & Liang, G. DNMT3B isoforms without catalytic activity stimulate gene body methylation as accessory proteins in somatic cells. Nat Commun 7, 11453, doi:10.1038/ncomms11453 (2016).

21 Weisenberger, D. J. et al. Role of the DNA methyltransferase variant DNMT3b3 in DNA methylation. Mol Cancer Res 2, 62–72 (2004).

22 Guo, X. et al. Structural insight into autoinhibition and histone H3-induced activation of DNMT3A. Nature 517, 640–644, doi:10.1038/nature13899 (2015).

23 Zhang, Y. et al. Chromatin methylation activity of Dnmt3a and Dnmt3a/3L is guided by interaction of the ADD domain with the histone H3 tail. Nucleic Acids Res 38, 4246–4253, doi:10.1093/nar/gkq147 (2010).

24 Weinberg, D. N. et al. The histone mark H3K36me2 recruits DNMT3A and shapes the intergenic DNA methylation landscape. Nature 573, 281–286, doi:10.1038/s41586-019-1534-3 (2019).

25 Dukatz, M. et al. H3K36me2/3 Binding and DNA Binding of the DNA Methyltransferase DNMT3A PWWP Domain Both Contribute to its Chromatin Interaction. J Mol Biol 431, 5063–5074, doi:10.1016/j.jmb.2019.09.006 (2019).

26 Qiu, C., Sawada, K., Zhang, X. & Cheng, X. The PWWP domain of mammalian DNA methyltransferase Dnmt3b defines a new family of DNA-binding folds. Nat Struct Biol 9, 217–224, doi:10.1038/nsb759 (2002).

27 Dhayalan, A. et al. The Dnmt3a PWWP domain reads histone 3 lysine 36 trimethylation and guides DNA methylation. J Biol Chem 285, 26114–26120, doi:10.1074/jbc.M109.089433 (2010).

28 Lu, J. et al. Structural basis for the allosteric regulation and dynamic assembly of DNMT3B. Nucleic Acids Res 51, 12476–12491, doi:10.1093/nar/gkad972 (2023).

29 Wapenaar, H. et al. The N-terminal region of DNMT3A engages the nucleosome surface to aid chromatin recruitment. EMBO Rep 25, 5743–5779, doi:10.1038/s44319-024-00306-3 (2024).

30 Lyons, D. B. & Zilberman, D. DDM1 and Lsh remodelers allow methylation of DNA wrapped in nucleosomes. Elife 6, doi:10.7554/eLife.30674 (2017).

31 Kelly, T. K. et al. Genome-wide mapping of nucleosome positioning and DNA methylation within individual DNA molecules. Genome Res 22, 2497–2506, doi:10.1101/gr.143008.112 (2012).

32 Voong, L. N. et al. Insights into Nucleosome Organization in Mouse Embryonic Stem Cells through Chemical Mapping. Cell 167, 1555–1570 e1515, doi:10.1016/j.cell.2016.10.049 (2016).

33 Valouev, A. et al. Determinants of nucleosome organization in primary human cells. Nature 474, 516–520, doi:10.1038/nature10002 (2011).

34 Brogaard, K., Xi, L., Wang, J. P. & Widom, J. A map of nucleosome positions in yeast at base-pair resolution. Nature 486, 496–501, doi:10.1038/nature11142 (2012).

35 Zhurkin, V. B. & Norouzi, D. Topological polymorphism of nucleosome fibers and folding of chromatin. Biophys J 120, 577–585, doi:10.1016/j.bpj.2021.01.008 (2021).

36 Gibson, B. A. et al. Organization of Chromatin by Intrinsic and Regulated Phase Separation. Cell 179, 470–484 e421, doi:10.1016/j.cell.2019.08.037 (2019).

37 Kelly, T. K. et al. H2A.Z maintenance during mitosis reveals nucleosome shifting on mitotically silenced genes. Mol Cell 39, 901–911, doi:10.1016/j.molcel.2010.08.026 (2010).

38 McGinty, R. K. & Tan, S. Principles of nucleosome recognition by chromatin factors and enzymes. Curr Opin Struct Biol 71, 16–26, doi:10.1016/j.sbi.2021.05.006 (2021).

39 Abini-Agbomson, S. et al. Catalytic and non-catalytic mechanisms of histone H4 lysine 20 methyltransferase SUV420H1. bioRxiv, doi:10.1101/2023.03.17.533220 (2023).

40 Wang, X. et al. Molecular analysis of PRC2 recruitment to DNA in chromatin and its inhibition by RNA. Nat Struct Mol Biol 24, 1028–1038, doi:10.1038/nsmb.3487 (2017).

41 Brohm, A. et al. Methylation of recombinant mononucleosomes by DNMT3A demonstrates efficient linker DNA methylation and a role of H3K36me3. Commun Biol 5, 192, doi:10.1038/s42003-022-03119-z (2022).

42 Ngo, T. T., Zhang, Q., Zhou, R., Yodh, J. G. & Ha, T. Asymmetric unwrapping of nucleosomes under tension directed by DNA local flexibility. Cell 160, 1135–1144, doi:10.1016/j.cell.2015.02.001 (2015).

43 Jumper, J. et al. Highly accurate protein structure prediction with AlphaFold. Nature 596, 583–589, doi:10.1038/s41586-021-03819-2 (2021).

44 Evans, R., et al. Protein complex prediction with AlphaFold-Multimer. bioRxiv, 2021.2010.2004.463034, doi:10.1101/2021.10.04.463034 (2022).

45 Wang, H., Farnung, L., Dienemann, C. & Cramer, P. Structure of H3K36-methylated nucleosome-PWWP complex reveals multivalent cross-gyre binding. Nat Struct Mol Biol 27, 8–13, doi:10.1038/s41594-019-0345-4 (2020).

46 Taglini, F. et al. DNMT3B PWWP mutations cause hypermethylation of heterochromatin. EMBO Rep 25, 1130–1155, doi:10.1038/s44319-024-00061-5 (2024).

47 Gretarsson, K. H. et al. Cancer-associated DNA hypermethylation of Polycomb targets requires DNMT3A dual recognition of histone H2AK119 ubiquitination and the nucleosome acidic patch. Sci Adv 10, eadp0975, doi:10.1126/sciadv.adp0975 (2024).

48 Chen, X. et al. Structural basis for the H2AK119ub1-specific DNMT3A-nucleosome interaction. Nat Commun 15, 6217, doi:10.1038/s41467-024-50526-3 (2024).

49 Poepsel, S., Kasinath, V. & Nogales, E. Cryo-EM structures of PRC2 simultaneously engaged with two functionally distinct nucleosomes. Nat Struct Mol Biol 25, 154–162, doi:10.1038/s41594-018-0023-y (2018).

50 Lowary, P. T. & Widom, J. New DNA sequence rules for high affinity binding to histone octamer and sequence-directed nucleosome positioning. J Mol Biol 276, 19–42, doi:10.1006/jmbi.1997.1494 (1998).

51 Dyer, P. N. et al. Reconstitution of nucleosome core particles from recombinant histones and DNA. Methods Enzymol 375, 23–44, doi:10.1016/s0076-6879(03)75002-2 (2004).

52 Kastner, B. et al. GraFix: sample preparation for single-particle electron cryomicroscopy. Nat Methods 5, 53–55, doi:10.1038/nmeth1139 (2008).

53 Stark, H. GraFix: stabilization of fragile macromolecular complexes for single particle cryo-EM. Methods Enzymol 481, 109–126, doi:10.1016/S0076-6879(10)81005-5 (2010).

54 Punjani, A., Rubinstein, J. L., Fleet, D. J. & Brubaker, M. A. cryoSPARC: algorithms for rapid unsupervised cryo-EM structure determination. Nat Methods 14, 290–296, doi:10.1038/nmeth.4169 (2017).

55 Sanchez-Garcia, R. et al. DeepEMhancer: a deep learning solution for cryo-EM volume post-processing. Commun Biol 4, 874, doi:10.1038/s42003-021-02399-1 (2021).

56 Wu, H. et al. Structural and histone binding ability characterizations of human PWWP domains. PLoS ONE 6, doi:10.1371/journal.pone.0018919 (2011).

57 Pettersen, E. F. et al. UCSF Chimera--a visualization system for exploratory research and analysis. J Comput Chem 25, 1605–1612, doi:10.1002/jcc.20084 (2004).

58 Adams, P. D. et al. PHENIX: a comprehensive Python-based system for macromolecular structure solution. Acta Crystallogr D Biol Crystallogr 66, 213–221, doi:10.1107/S0907444909052925 (2010).

59 Emsley, P. & Cowtan, K. Coot: model-building tools for molecular graphics. Acta Crystallogr D Biol Crystallogr 60, 2126–2132, doi:10.1107/S0907444904019158 (2004).

