## Supplementary Figures for "The structural basis for *de novo* DNA methylation in chromatin"

Figure S1

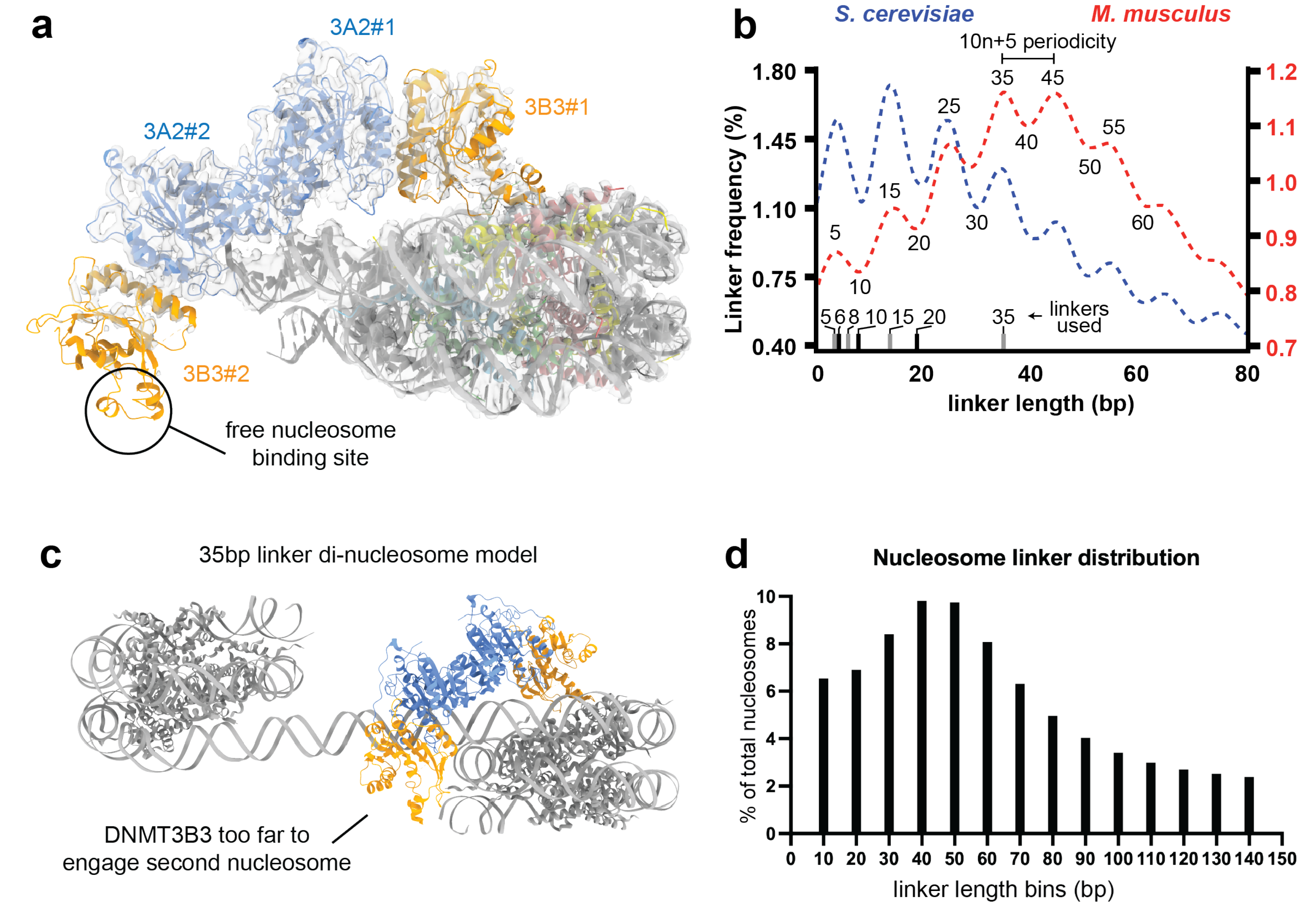


**Figure S1: constraints of di-nucleosome interactions with the DNMT3A2/3B3 tetramer**
**a** Cryo-EM structure and model (PDB:6PA7, EMDB: 20281) of the DNMT3A2/3B3 tetramer bound to a di-nucleosome to show the unsatisfied nucleosome binding site on DNMT3B3 #2. **b** Linker length distributions for S. cerevisiae (ref 34) and M musculus (ref 32), based on ref 36. **c** Manually generated structural model of a di-nucleosome separated 35bp linker. The structure of DNMT3A2/3B3 bound to a mono-nucleosome (PDB 6PA7, Cyan) is superimposed onto one of the two nucleosomes to show the position of the DNMT tetramer during linker methylation. **d** All nucleosomes linker lengths in mouse ES cells (ref 32) separated into 10bp bins.

Figure S2

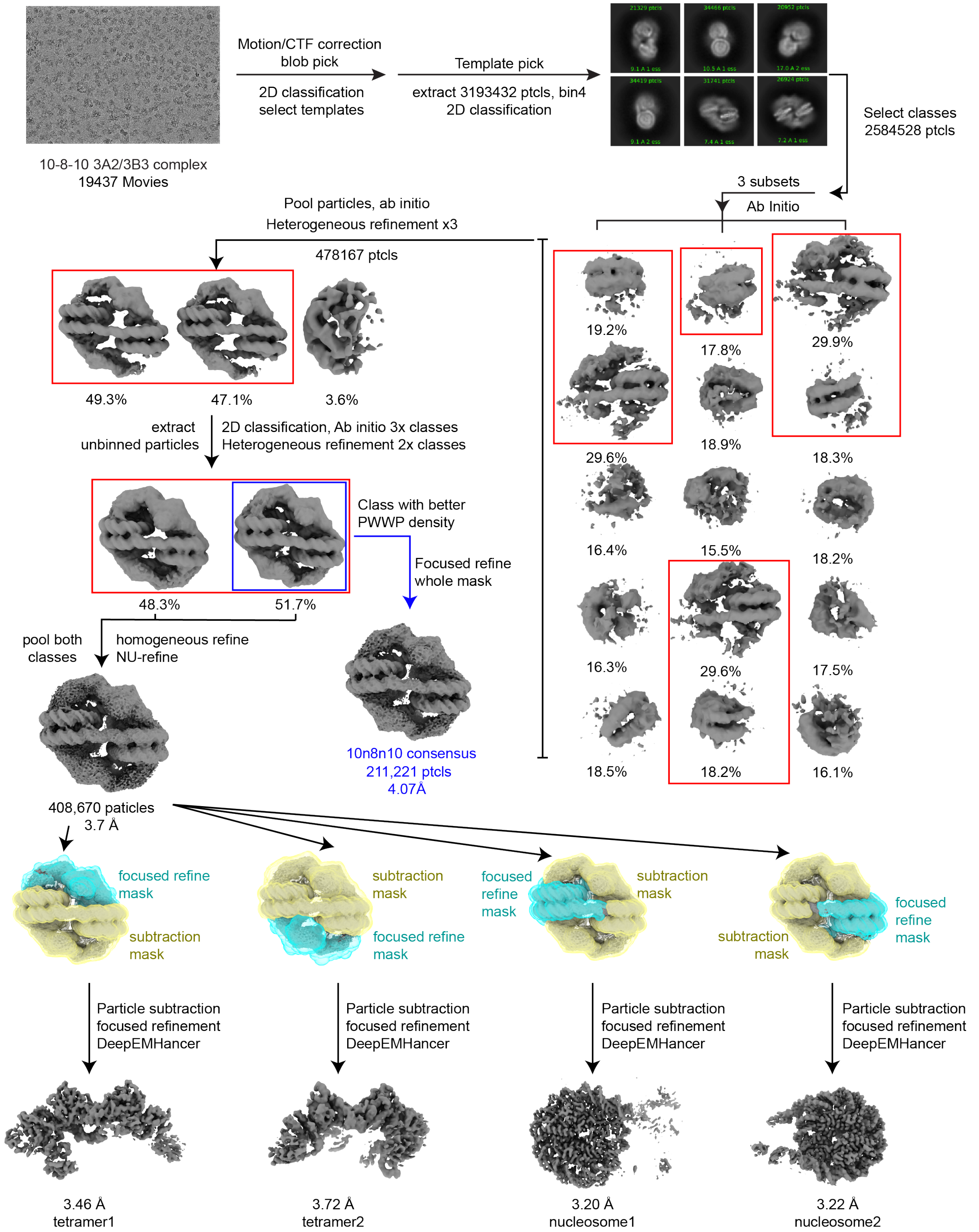


**Figure S2: Cryo-EM processing pipeline for the complex between DNMT3A2/3B3 and the 10-8-10 di-nucleosome**

Figure S3

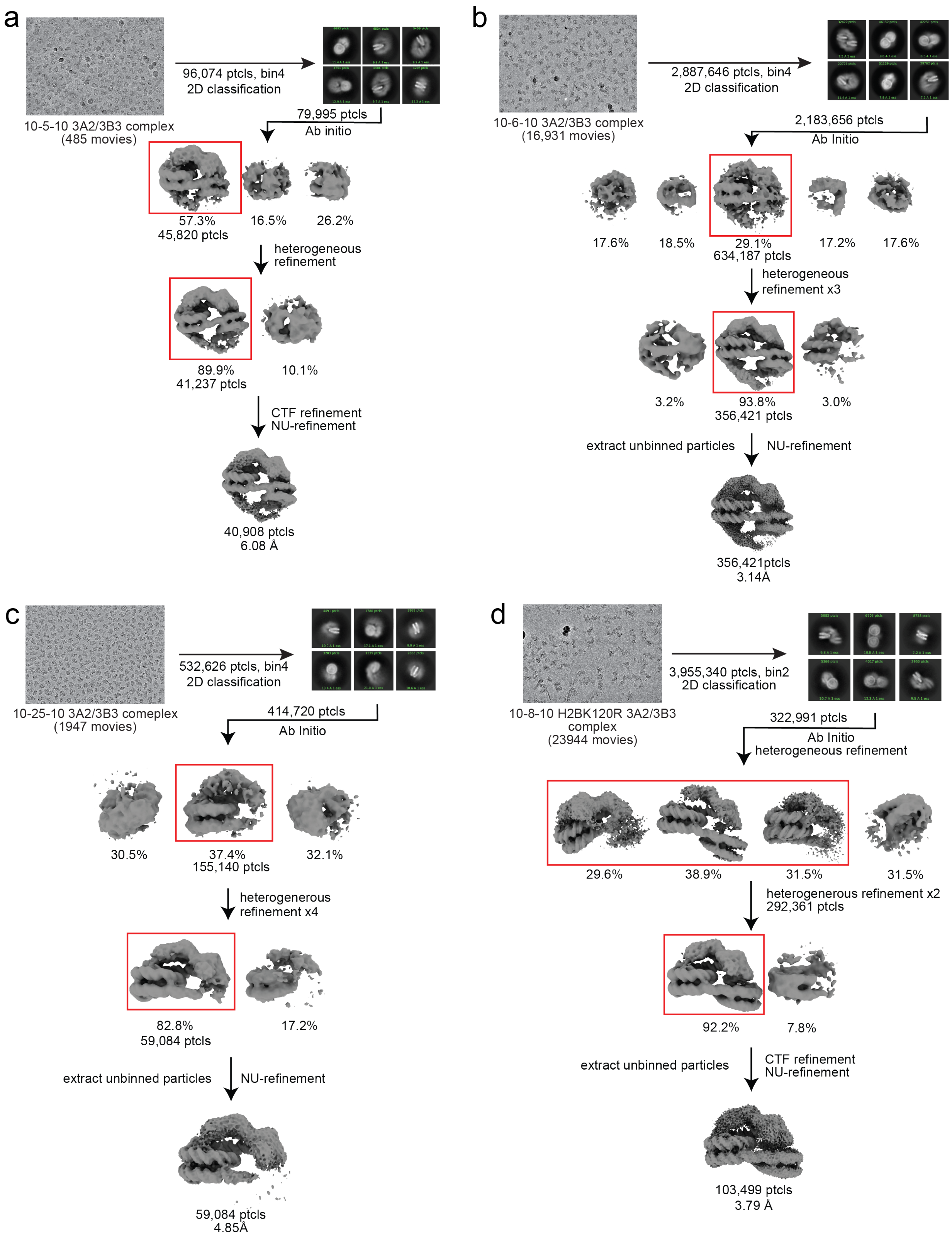


**Figure S3: Cryo-EM processing pipeline for the complex between DNMT3A2/3B3 and the 10-5-10 di-nucleosome (a), 10-6-10 di-nucleosome (b), 10-25-10 di-nucleosome (c) and the 10-8-10 H2BK120R di-nucleosome (d).**

Figure S4

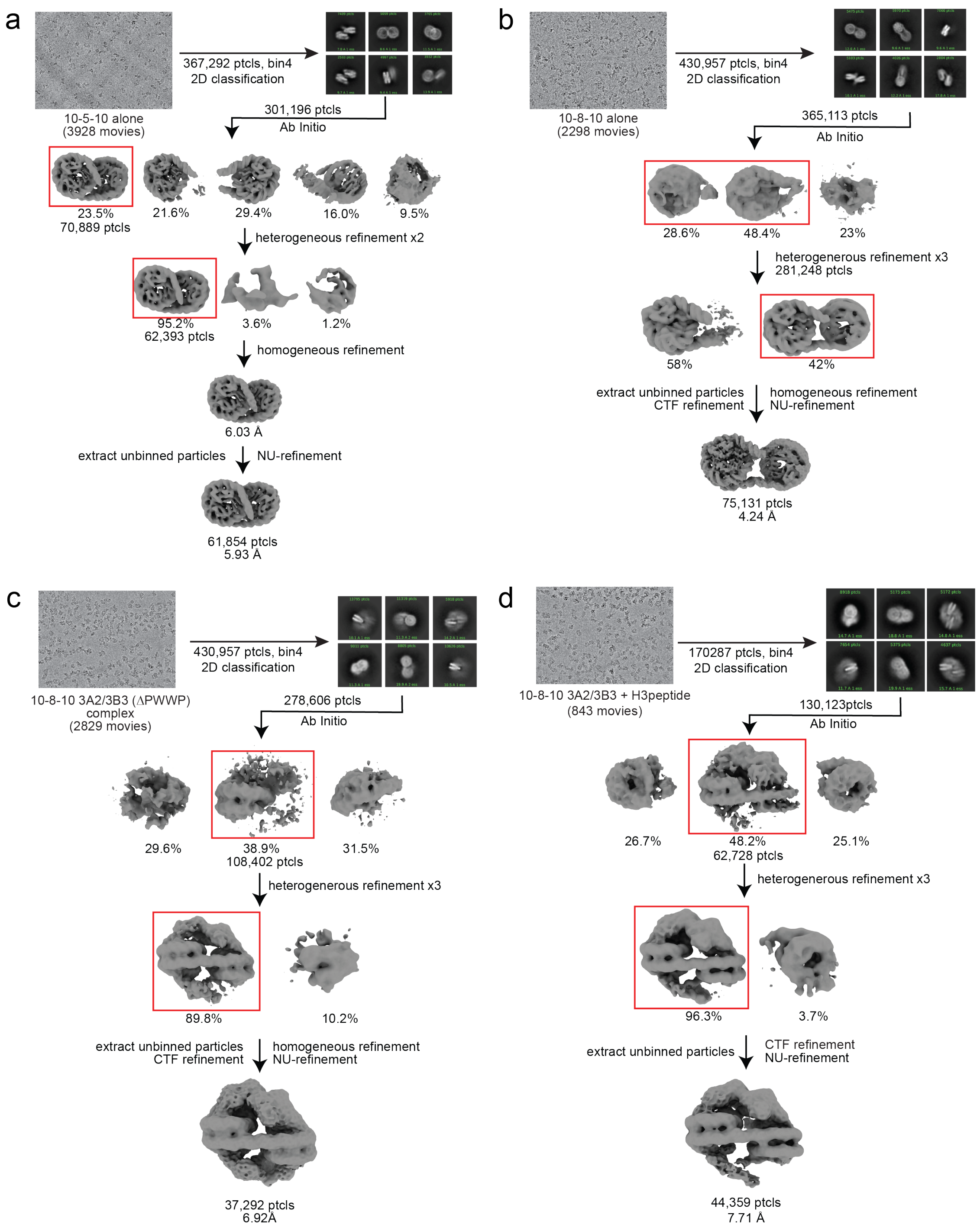


**Figure S4: Cryo-EM processing pipeline for the 10-5-10 di-nucleosome alone (a), the 10-8-10 di-nucleosome alone (b), the 10-8-10 di-nucleosome DNMT3A2/3B3 (ΔPWWP) complex (c) and the 10-8-10 di-nucleosome DNMT3A2/3B3 complex in excess H3 peptide (d)**

Figure S5

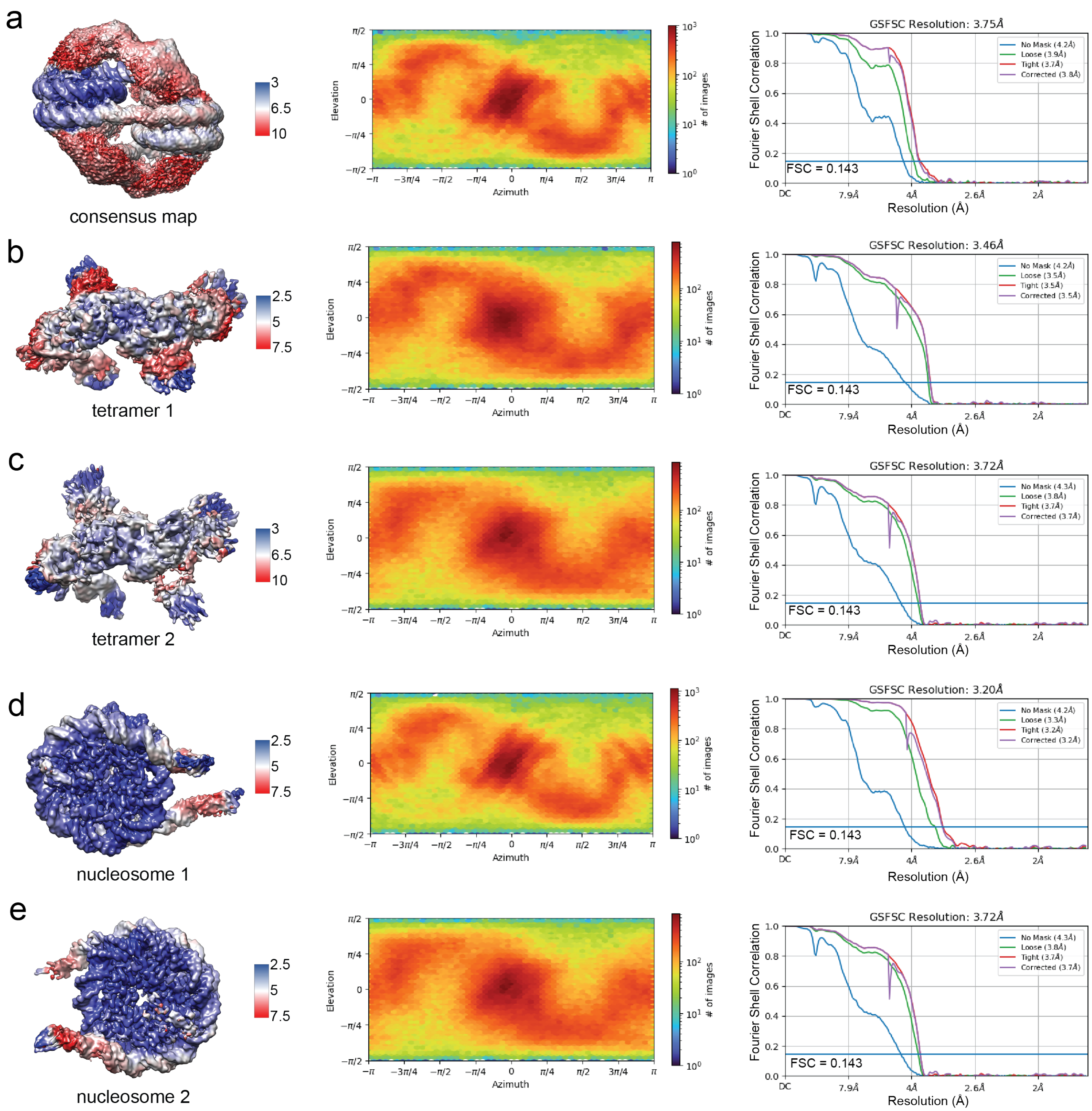


**Figure S5: Cryo-EM map validation for the 10-8-10 di-nucleosome complex with DNMT3A2/3B3.**
Local resolution estimates (left), particle orientation distribution (middle), and FSC curves (right) for, **a** the 10-8-10 di-nucleosome complex and for the focused maps of **b** tetramer 1, **c** tetramer 2, **d** nucleosome 1 and **e** nucleosome 2.

Figure S6

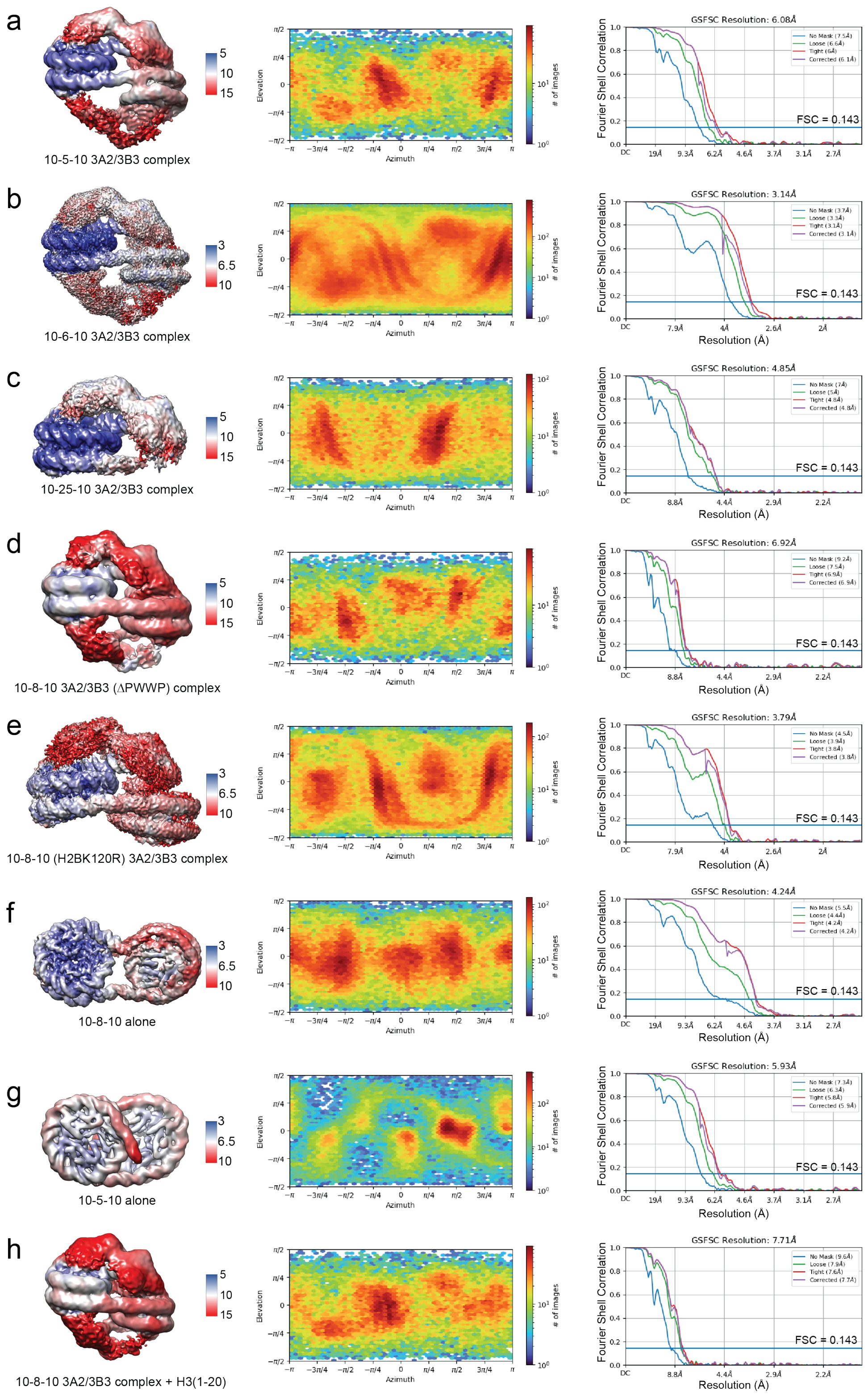


**Figure S6: Cryo-EM map validation for di-nucleosome complexes.**
Local resolution estimates (left), particle orientation distribution (middle), and FSC curves (right) for the DNMT3A2/3B3 10-5-10 di-nucleosome complex (**a**) DNMT3A2/3B3 10-6-10 di-nucleosome complex (**b**), DNMT3A2/3B3 10-25-10 di-nucleosome complex (**c**), DNMT3A2/3B3 (ΔPWWP) 10-8-10 di-nucleosome complex (**d**), DNMT3A2/3B3 10-8-10 (H2BK120R) di-nucleosome complex (**e**), 10-8-10 di-nucleosome alone (**f**) 10-5-10 di-nucleosome alone (**g**) and the DNMT3A2/3B3 10-8-10 di-nucleosome complex in the presence of excess H3(1-20) peptide (**h**)

Figure S7

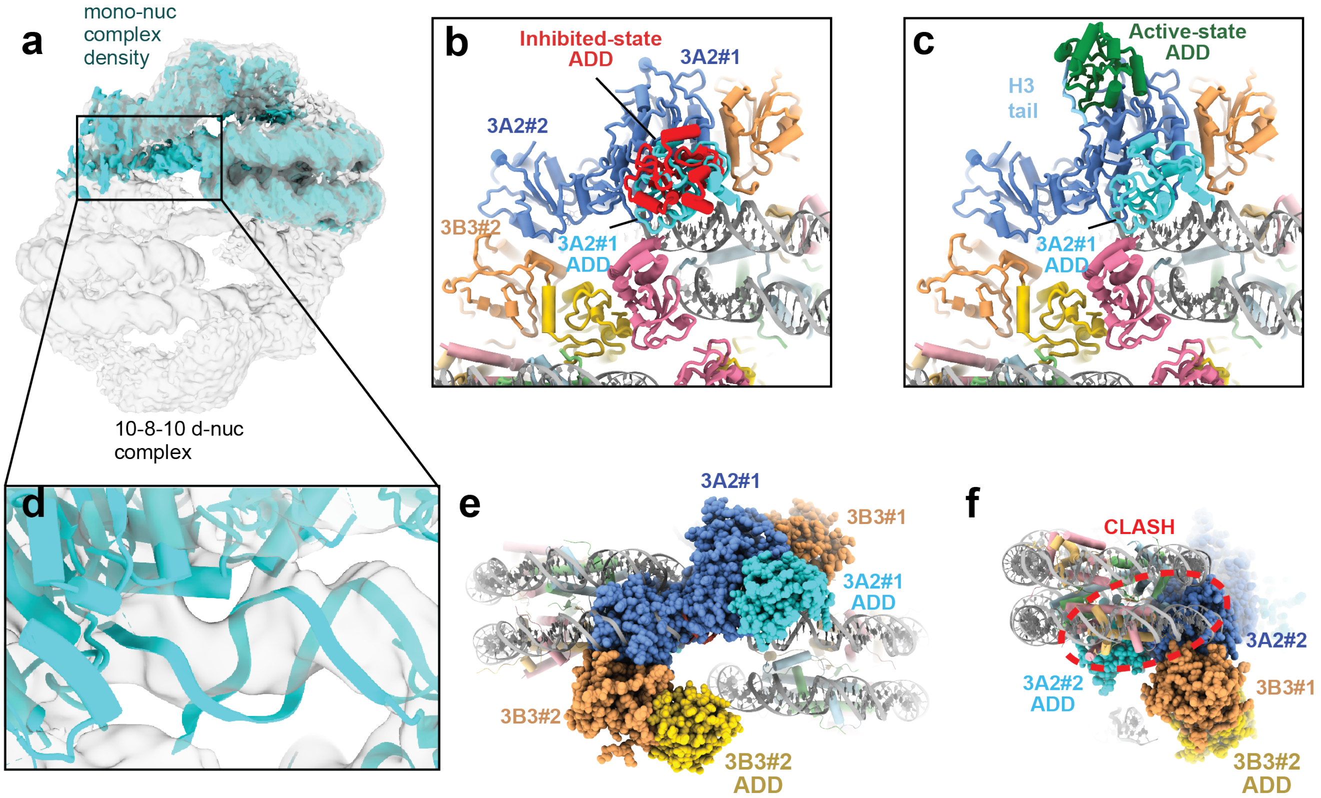


**Figure S7: Details of di-nucleosome binding by DNMT3A2/3B3**

**a** EM density for the mono-nucleosome bound structure of DNMT3A2/3B3 (EMDB: 20281, cyan) superimposed with the density of the 10-8-10 di-nucleosome DNMT3A2/3B3 complex, depicted as a semitransparent gray surface. **b** structural model of the 10-8-10 di-nucleosome DNMT3A2/3B3 complex where DNMT3A2 is superimposed with the structure of inhibited-state DNMT3A1 (PDB: 4U7P) to show the relative location of the ADD domains. **c** Same as in **b** except that DNMT3A2 is superimposed with the structure of active-state DNMT3A1 to show the ADD conformation change. **d** structural model of the mono-nucleosome bound of DNMT3A2/3B3 (PDB 6PA7, Cyan) fit into the density of the 10-8-10 di-nucleosome DNMT3A2/3B3 complex. **e-f** Structural model of the di-nucleosome-bound DNMT3A2/3B3 tetramer oriented for linker methylation of the 10-8-10 di-nucleosome. The close spacing of the two nucleosomes results in a strong steric clash between DNMT3A2#2 and the second nucleosome.

Figure S8

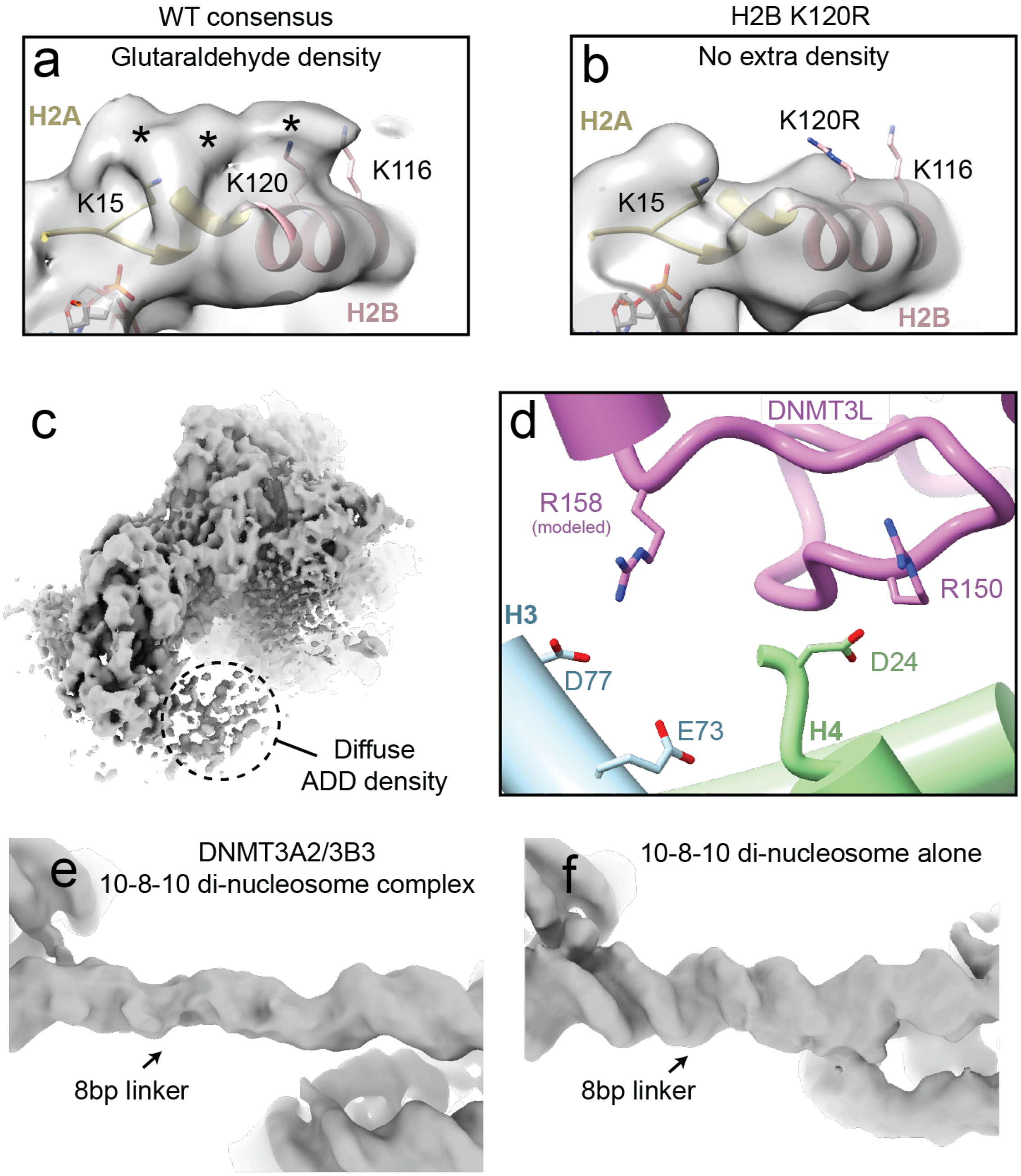


**Figure S8: DNMT interactions with the histone core.**
**a** Close up view of the extra glutaraldehyde density near the acidic patch in the consensus map of the 10-8-10 di-nucleosome DNMT3A2/3B3 complex. Extra density is indicated using *. **b** Close up of the same region of the H2BK120R 10-8-10 di-nucleosome in complex with DNMT3A2/3B3. No density for glutaraldehyde crosslinks is observed the H2BK120R mutant nucleosome. **c** Cryo-EM density for the DNMT3A2/3B3 hetero-tetramer bound to a mono-nucleosome (EMDB: 20281) showing diffuse density for the DNMT3B3 ADD domain. **d** The ADD domain of DNMT3L (PDB: 2PV0) superimposed with the DNMT3B3 #2 ADD domain (not shown) showing the location of positively charged residues near negative charged residues in H3 and H4. The sidechain of R158 was not included in PDB: 2PV0, so a high probability rotamer for R158 is modeled. **e** Cryo-EM density for the DNA linker in the 10-8-10 di-nucleosome, DNMT3A2/3B3 complex. **f** Cryo-EM density for the DNA linker in the 10-8-10 di-nucleosome alone.

Figure S9

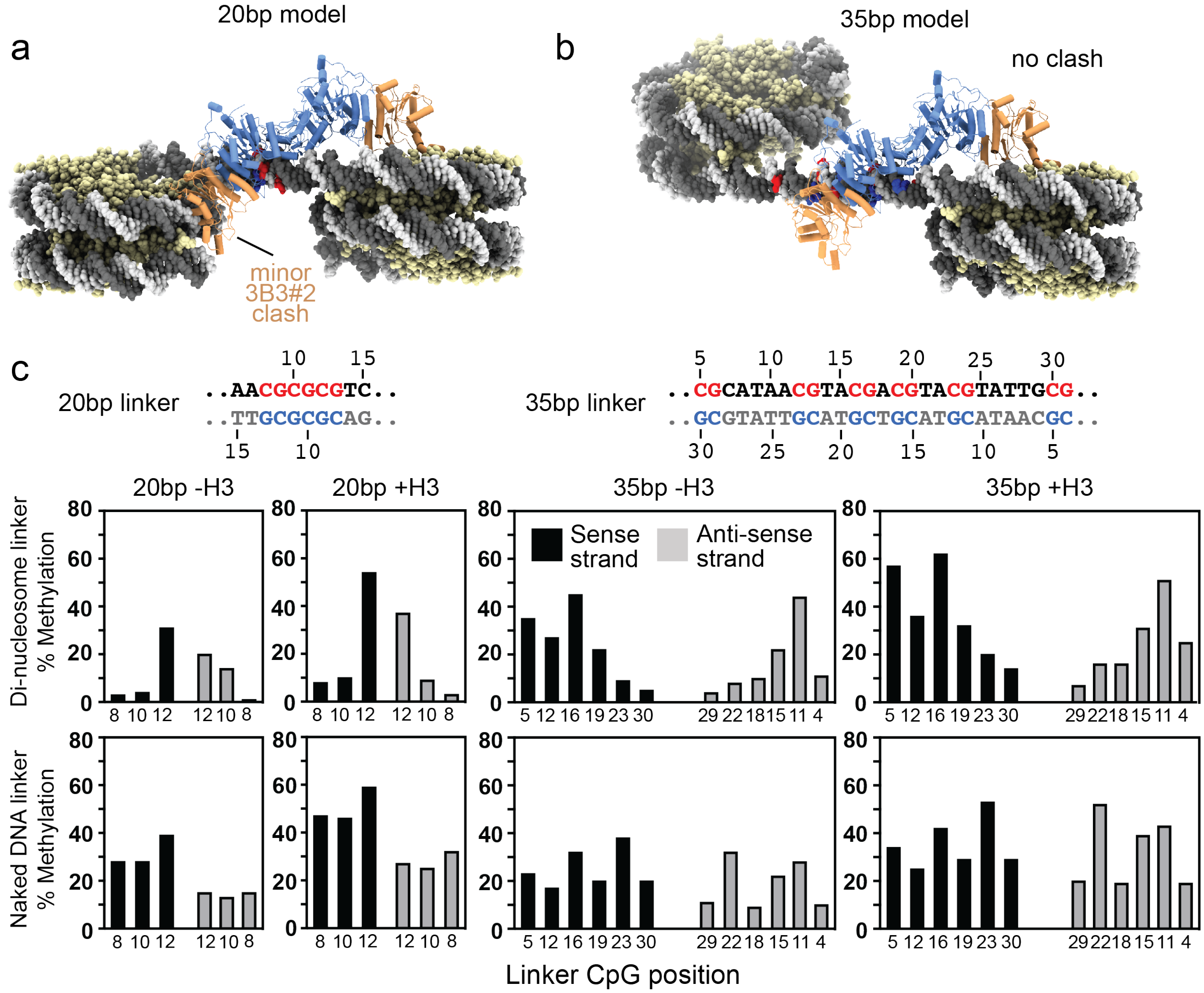


**Figure S9: DNMT3A2/3B3 linker methylation of the 20bp and 35bp di-nucleosome**
**a-b** Manually generated model of the 20bp (a) and 35bp (b) di-nucleosomes superimposed with the cryo-EM structure of the DNMT3A2/3B3 tetramer bound to a mono-nucleosome (PDB: 6PA7). **c** Bisulfite sequencing assays showing the percentage of cytosine methylation at each CpG position for the 20bp and 30bp di-nucleosome with and without excess H3 (1-20) peptide in the reaction.

Figure S10

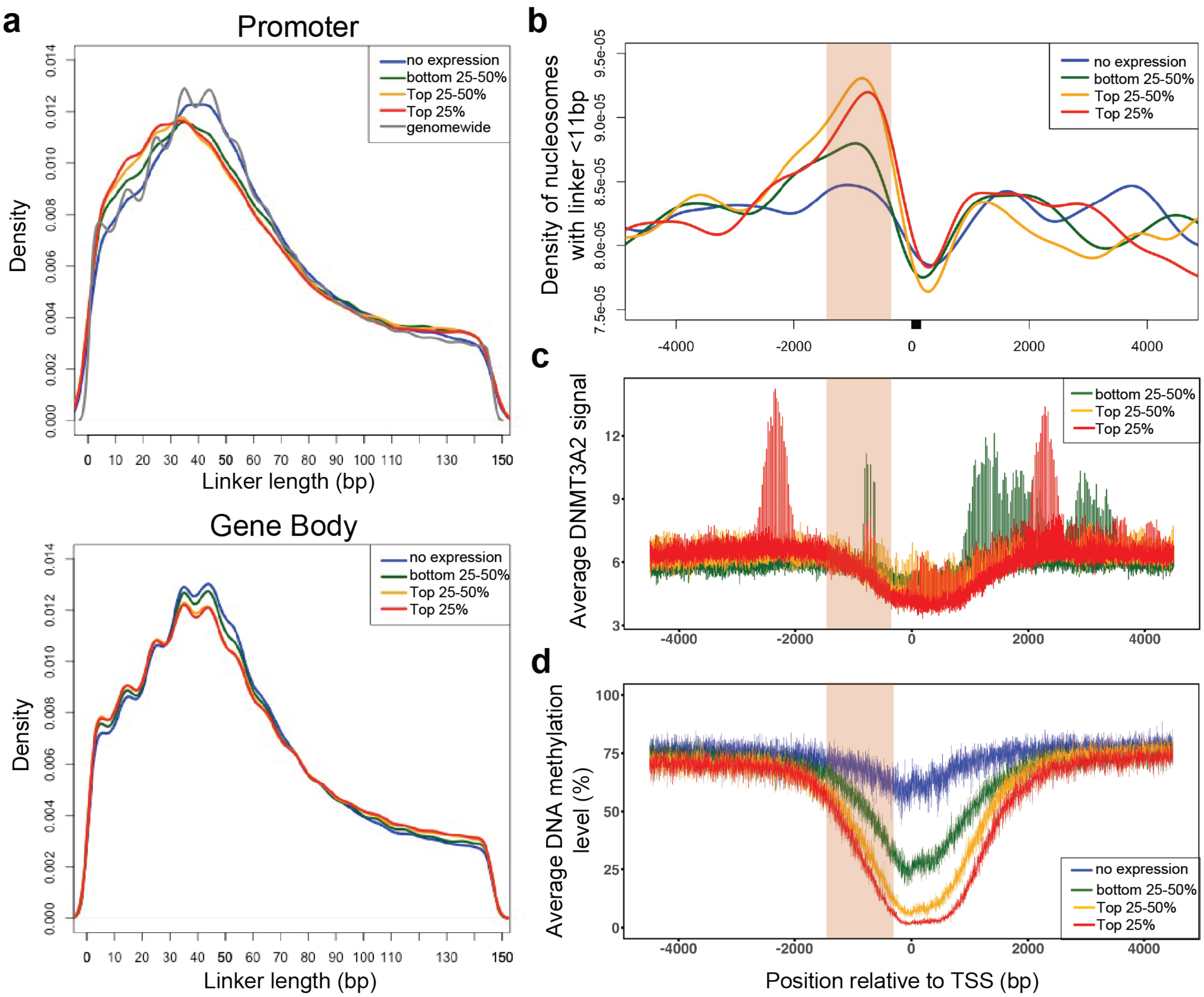


**Figure S10: Linker length shortening at active promotor regions**
**a** Nucleosome linker length distribution in gene promoter (±2000bp relative to transcription start site (TSS), top panel) and gene body (bottom panel) by gene expression quartiles in FPKM shows higher expressed genes are enriched with shorter linker lengths especially at promoters. **b** Distribution of nucleosomes with linker shorter than 11 bp around TSS shows that these nucleosomes are enriched at gene promoters, with peaks around 1000bp upstream of TSS for higher expressed genes. **c** Average DNMT3A2 signal (GSM3772691) around TSS shows a peak for DNMT3A2 binding that colocalize with the peak of nucleosomes with short linkers. **d** Average DNA methylation level around TSS shows that the peak of nucleosomes with short linkers are in the border between the methylated and unmethylated regions upstream of TSS.

Figure S11

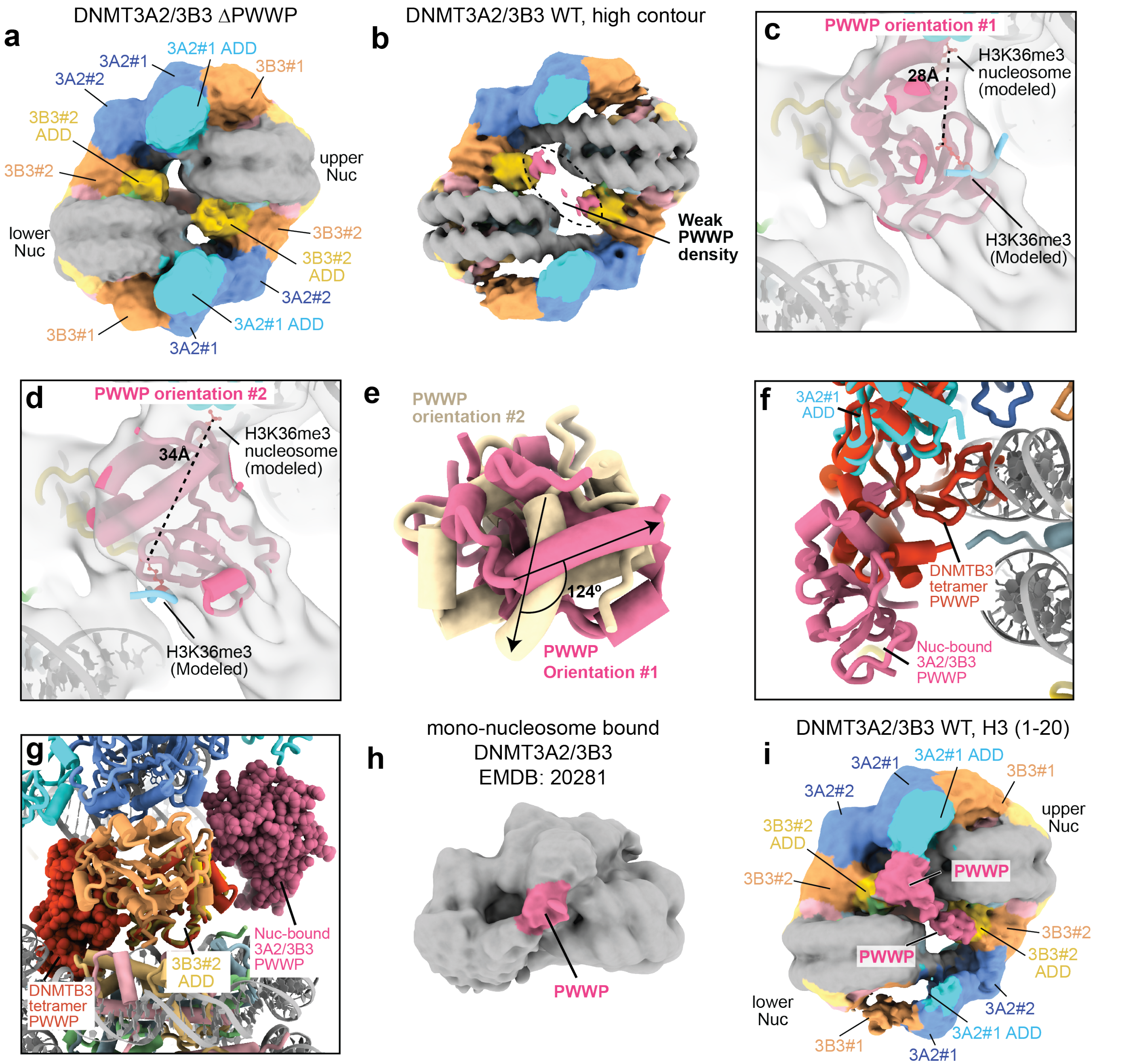


**Figure S11: Details of PWWP docking**
**a** Cryo-EM map for the 10-8-10 di-nucleosome in complex with the DNMT3A2(476-912)/3B3(Δ234-286) variant that lacks the PWWP domains of DNMT3A2 and DNMT3B3 (ΔPWWP). No density for the PWWP domains is evident in the map. **b** Consensus Cryo-EM map for the 10-8-10 di-nucleosome filtered to 10Å and displayed at high contour. **c-d** Two different conformations of PWWP domain docked into consensus Cryo-EM density for the 10-8-10 di-nucleosome in complex with WT DNMT3A2/3B3 filtered to 10Å and displayed as a semi-transparent gray surface. **e** comparison of two docking orientations showing that they are rotated relative to each other. **f-g** comparison between the orientation of the docked PWWP domain and the structure of the PWWP-ADD interaction in the DNMT3B1 homotetramer (PDB: 8EIH). **f** The position of the DNMT3B1 PWWP domain (red) is determined by superimposing the DNMT3B1 ADD domain with the ADD domain of DNMT3A2 #1. **g** The position of the DNMT3B1 PWWP domain (red spheres) is determined by superimposing the DNMT3B1 ADD domain with the ADD domain of DNMT3B3 #2. **h** Cryo-EM map of the DNMT3A2/3B3 mono-nucleosome complex (EMDB: 20281) gaussian filtered with a standard deviation of 2.8 in ChimeraX. Density for the PWWP domain is colored pink. **i** Cryo-EM map of the 10-8-10 di-nucleosome in complex with WT DNMT3A2/3B3 determined in the presence of excess unmodified H3(1-20) peptide.
